# Engineering sequestration-based biomolecular classifiers with shared resources

**DOI:** 10.1101/2024.04.15.589451

**Authors:** Hossein Moghimianavval, Ignacio Gispert, Santiago R. Castillo, Olaf B. W. H. Corning, Allen P. Liu, Christian Cuba Samaniego

## Abstract

Constructing molecular classifiers that enable cells to recognize linear and non-linear input patterns would expand the biocomputational capabilities of engineered cells, thereby unlocking their potential in diagnostics and therapeutic applications. While several biomolecular classifier schemes have been designed, the effect of biological constraints such as resource limitation and competitive binding on the function of those classifiers has been left unexplored. Here, we first demonstrate the design of a sigma factor-based perceptron as a molecular classifier working on the principles of molecular sequestration between the sigma factor and its anti-sigma molecule. We then investigate how the output of the biomolecular perceptron, *i.e*., its response pattern or decision boundary, is affected by the competitive binding of sigma factors to a pool of shared and limited resources of core RNA polymerase. Finally, we reveal the influence of sharing limited resources on multi-layer perceptron neural networks and outline design principles that enable the construction of non-linear classifiers using sigma-based biomolecular neural networks in the presence of competitive resource-sharing effects.

## 1 Introduction

Cellular biocomputation is prevalent in nature with examples including activation of genetic circuits during cell proliferation, decision-making in immune response, and a myriad of phosphorylation-based signaling pathways for determining correct response to exogenous signals [1, 2, 3]. The foundation of such computational processes is often laid on molecular interactions such as protein dimerization or ligand-receptor binding. Thus, the inputs of the computational modules in biological systems are typically the concentration of certain monomeric molecules or ligands and, similarly, the outputs are the concentration of specific dimeric or multimeric molecules.

Drawing inspiration from natural systems, synthetic biologists have been striving to engineer biocomputational schemes in top-down as well as bottom-up synthetic biological systems. While the majority of biocomputational designs rely on utilizing genetic circuits to engineer basic logic gates and simple computational tasks [4, 5, 6], a few studies have demonstrated engineering protein-based circuits that utilize proteolytic or phosphorylation reactions to generate an output [7, 8, 9]. Although such biocomputational modules enable simple tasks such as biosensing of chemical species and basic computation, they typically generate digital (0 or 1 or “on or off”) responses. Furthermore, encoding more sophisticated processing using logic gates demands intricate architectures with many logic gates and computational layers, rendering them convoluted for practical applications for complex tasks.

Therefore, constructing simple signal processing units inside living systems that can perform intricate computational tasks such as classification and decision-making are of great interest (**Fig. 1A**, left). Implementing molecular classifiers in living cells would enable the creation of ultra-sensitive biosensors, programming accurate cellular responses through molecular circuits, and enhanced discrimination of inputs through combinatorial sensing [10]. For example, a simple linear classifier (**Fig. 1A**, middle) equips a cell with a signal processing system that ideally allows output generation only in certain input regimes (where *x*_1_ and *x*_2_ approach 1 in the example). Further, combining different molecular classifiers results in more complex, non-linear computation, thus expanding the capabilities of cellular biocomputation.

**Figure 1:**
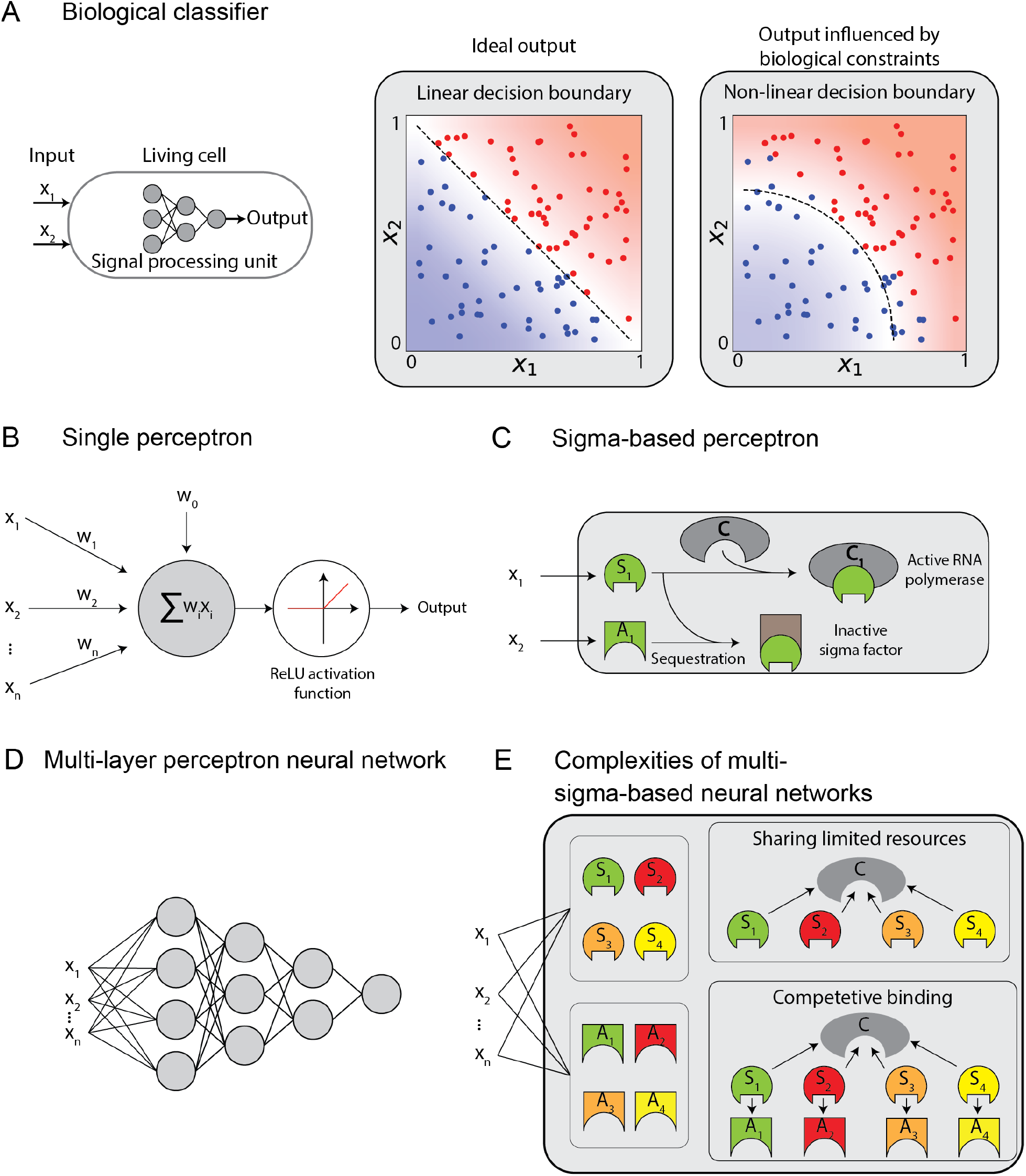
Designing a Biomolecular Neural Network (BNN) utilizing a sigma-based perceptron and multilayer neural networks with shared limited resources for molecular classification: **A**: A molecular classifier as a signal processing unit in living cells enables more sophisticated biocomputation. The resource constraints in biological systems, however, may perturb the designed decision boundary. **B**: The abstract representation of a perceptron, the building block of deep neural networks. **C**: The representation of the biological perceptron design in a sigma-based system using sequestration of a sigma factor and its corresponding anti-sigma protein. The inputs of the perceptron are concentrations of the sigma and anti-sigma. **D**: Abstract depiction of multi-layer neural networks made of multiple perceptrons in each layer. **E**: Schematic design of a multi-layer perceptron in a sigma-based system that poses two major limitations: sharing limited resources and competitive binding. These complexities influence the decision boundary of the multi-layer neural network.

Binary classification using a linear decision boundary was demonstrated in the field of artificial intelligence (AI) in 1958 [11]. A simple computational unit called ‘perceptron’ performs binary classification by computing the linear combination of weighted inputs and passing the summed weighted inputs through an activation function. The most popular activation function in modern perceptrons is a thresholding function called Rectified Linear Unit known as ReLU, in which the output is larger than zero when the input crosses a threshold (**Fig. 1B)**. The collective processing of inputs by many layers of multiple perceptrons, known as deep neural networks or artificial neural networks (ANNs), can result in the recapitulation of any continuous function [12], thus making ANNs capable of performing complicated tasks such as non-linear classification and accurate prediction [13, 14, 15, 16].

The simple architecture of a perceptron has motivated many efforts towards the creation of a biological perceptron as the signal processing unit for linear classification. The construction of a single biological perceptron can also pave the way for implementing non-linear input classification in living cells by utilizing multiple perceptrons, thus creating biomolecular neural networks (BNNs) (**Fig. 1D**). A biological perceptron must demonstrate the principal characteristics of an ANN (or computer-based) perceptron, *i.e*., the biological perceptron must include controllable molecular elements that determine the perceptron’s input weights and activation function.

However, as opposed to ANN perceptrons, biological perceptrons face challenges in linear classification due to biological constraints such as limited resources, competitive binding, and non-specific binding between molecules. Resource constraints in biological systems have been shown to significantly impact the function of biocomputational modules based on molecular interactions. Competitive binding of only a few promiscuous ligands to various receptors has been demonstrated to accommodate a wide range of signaling activities in multicellular organisms [17]. Similarly, it was recently shown that competitive protein dimerization allows small protein monomer networks to encode an extensive range of homo- or hetero-dimeric outputs through precise adjustments in the concentration of network monomers [18]. In addition, competing transcription factors that bind to the same RNA polymerase may affect the output of engineered genetic circuits in bacteria. [19, 20].These resource constraints can cause perturbations to the biological perceptron function, thus influencing its decision boundary (**Fig. 1A**, right). Subsequently, BNNs made from the biological perceptron with an altered output will also generate decision boundaries that may not closely follow their ideal design.

A few biological perceptrons have been demonstrated by using inducible gene expression networks [21, 22, 23, 24], enzymatic processing of different metabolites[25], principles of DNA strand-displacement [26, 27, 28, 29], and DNA-processing enzymes [30]. Relying on the sequestration of two interacting proteins, a biomolecular classifier with tunable positive and negative weights was designed computationally [31] and tested experimentally to achieve non-linear classification in mammalian cells [24]. Similarly, a phosphorylation-based neural network with positive and negative weights that perform non-linear classification (*i.e*., recapitulating XNOR and XOR) was designed by Cuba Samaniego *et al*. [32] Recently, a protein-based neural network that achieves linear classification was implemented by exploiting coiled-coil dimerization of engineered peptides [33]. Although these studies utilize different approaches to create a biological perceptron, they all rely on competitive interactions between input-processing molecules with a shared pool of limited resources. However, the effects of such resource constrains on the function of these perceptrons have remained unexplored.

Here, we develop a mathematical model to simulate a biological perceptron based on sigma factors, transcription factors that bacteria naturally use to control gene expression. Leveraging competitive dimerization between a sigma and either its corresponding anti-sigma molecule or RNA polymerase, we demonstrate the design of a simple perceptron with a non-linear activation function capable of realizing positive and negative weights (**Fig. 1C)**. We then impose two physiological requirements on our model to account for both competition between the input sigma factor and other present sigma factors as well as the limited available resources (*i.e*., core RNA polymerase). We show that resource sharing reveals its effect on the function of the perceptron by suppressing the output while introducing a slight perturbation to the linear decision boundary. Lastly, since engineering non-linear decision boundaries require multilayer perceptron networks (as depicted in **Fig. 1D**), we explore designing sequestration-reliant multi-layer sigma-based perceptron networks in the presence of perturbations caused by sharing limited resources (**Fig. 1E**). We demonstrate that resource sharing leads to deviations from ideal design that affect the output of the multi-layer perceptron network. However, despite the non-ideal function of perceptrons due to resource sharing and limited resources, we outline simple design principles for encoding non-linear response patterns such that they closely resemble their ideal design. Our analyses of biological perceptron and BNN function in the presence of resource constraints can also be utilized to model molecular classifiers *in silico* for more precise prediction or design of outputs of these biocomputational systems.

Since sharing limited resources is not exclusive to the sigma-based neural networks, our findings can be extended to other biocomputational input-processing systems used in living cells. Taking Clustered Regularly Interspaced Short Palindromic Repeats (CRISPR) gene editing as an example, the sequestration of a single guide RNA (sgRNA) strand by its complementary RNA, or anti-guide RNA, is analogous to the interaction of a sigma factor with its anti-sigma protein. If the sgRNA is engineered to drive a CRISPR reaction, the Cas protein will consequently be the limited resource that all sgRNAs will compete to bind to. Similarly, a limited pool of proteolytic substrates that are catalyzed by a protease-based neural network [34, 35] will follow the functional principles outlined in this work. Therefore, the principles of input classification under the effects of sharing limited resources outlined in this paper can be extended and used for designing other molecular classifiers.

## 2 Results

Throughout the manuscript, we indicate chemical species with capital letters (*e.g*., *X*) and their concentration with the corresponding lowercase letters (*e.g*., *x*). Table 1 summarizes all parameters used in the manuscript for mathematical analyses and computational simulations.

**Table 1:** List of parameters used in computational simulations. The value of parameters indicated with * are described in their corresponding figures.

| Parameter | Symbol | Unit | Value | Figure |
| --- | --- | --- | --- | --- |
| Perceptron input species | $[x_1, x_2]$ | $\mu M$ | $[0 \rightarrow 1]$ | 2,3,4 |
| Competing perceptron input species | $[\alpha, \beta]$ | $\mu M$ | $[0, 0.5, 1]$ | 2 |
| Sigma factor | $s_1, s_2$ | $\mu M$ | variable | 2,3,4 |
| Anti-sigma factor | $a_1, a_2$ | $\mu M$ | variable | 2,3,4 |
| Core RNA polymerase (RNAPol) | $c$ | $\mu M$ | $[0 \rightarrow 1]$ | 2,3,4 |
| Sigma $i$-RNAPol complex | $c_i$ | $\mu M$ | variable | 2,3,4 |
| Sigma-anti-sigma sequestration rate | $\gamma_1$ | $\mu M^{-1} h^{-1}$ | * | 2 |
| Sigma-RNAPol complex formation rate | $\gamma_2$ | $\mu M^{-1} h^{-1}$ | 10 | 2 |
| Competitive binding ratio | $r$ | - | * | 2,3,4 |
| Degradation rate | $\delta$ | $\mu M^{-1} h^{-1}$ | 1 | 2,3,4 |
| Node 1 input weights | $[w_1, w_2]$ | $h^{-1}$ | $[1, 1]$ | 3 |
| Node 2 input weights | $[\alpha_2, \beta_2]$ | $h^{-1}$ | $[0, 0]$ or $[0.5, 0.5]$ | 3 |
| Node 1 input weights | $[w_0^1, w_1^1, w_2^1]$ | $h^{-1}$ | $[1, 1, 1]$ | 4A |
| Node 2 input weights | $[-w_0^2, -w_1^2, w_2^2]$ | $h^{-1}$ | $[0.5, 1, 1]$ | 4A |
| Node 1 input weights | $[-w_0^1, w_1^1, w_2^1]$ | $h^{-1}$ | $[1.2, 1, 1]$ | 4E |
| Node 2 input weights | $[w_0^2, -w_1^2, -w_2^2]$ | $h^{-1}$ | $[0.7, 1, 1]$ | 4E |
| Node 3 input weights | $[-w_0^3, w_1^3, w_2^3]$ | $h^{-1}$ | $[0.15, 4, 4]$ | 4E |
| Node 1 input weights | $[w_0^1, -w_1^1, -w_2^1]$ | $h^{-1}$ | $[0.4, 0.5, 0.5]$ | 4H |
| Node 2 input weights | $[-w_0^2, w_1^2, -w_2^2]$ | $h^{-1}$ | $[0.8, 1.5, 0.5]$ | 4H |
| Node 3 input weights | $[-w_0^3, -w_1^3, w_2^3]$ | $h^{-1}$ | $[0.8, 0.5, 1.5]$ | 4H |
| Node 4 input weights | $[w_0^4, -w_1^4, -w_2^4, -w_3^4]$ | $h^{-1}$ | $[0.3, 1, 1, 1]$ | 4H |

### 2.1 Design of a rectified linear activation unit (ReLU) based on sigma factor-anti sigma factor interaction in the presence of shared limited resources

Molecular sequestration is the stoichiometric binding between two species that results in the formation of a dimeric complex. An example of molecular sequestration is the interaction between sigma factors and their corresponding anti-sigma proteins that leads to the formation of a complex that is unable to promote gene expression (**Fig. 1B**). Such interaction can be modeled assuming the sigma factor *S*_1_ and anti-sigma factor *A*_1_ are produced from species *X*_1_ and *X*_2_ at rate constants *w*_1_ and *w*_2_, respectively. Additionally, the produced proteins *S*_1_ and *A*_1_ degrade at rate *δ* and the sequestration occurs with rate constant *γ*_1_ (as shown in **Fig. 2A**). We summarize the list of chemical reactions as follows:

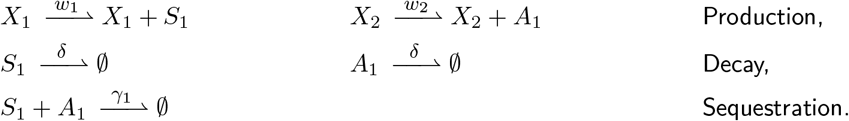

**Figure 2:**
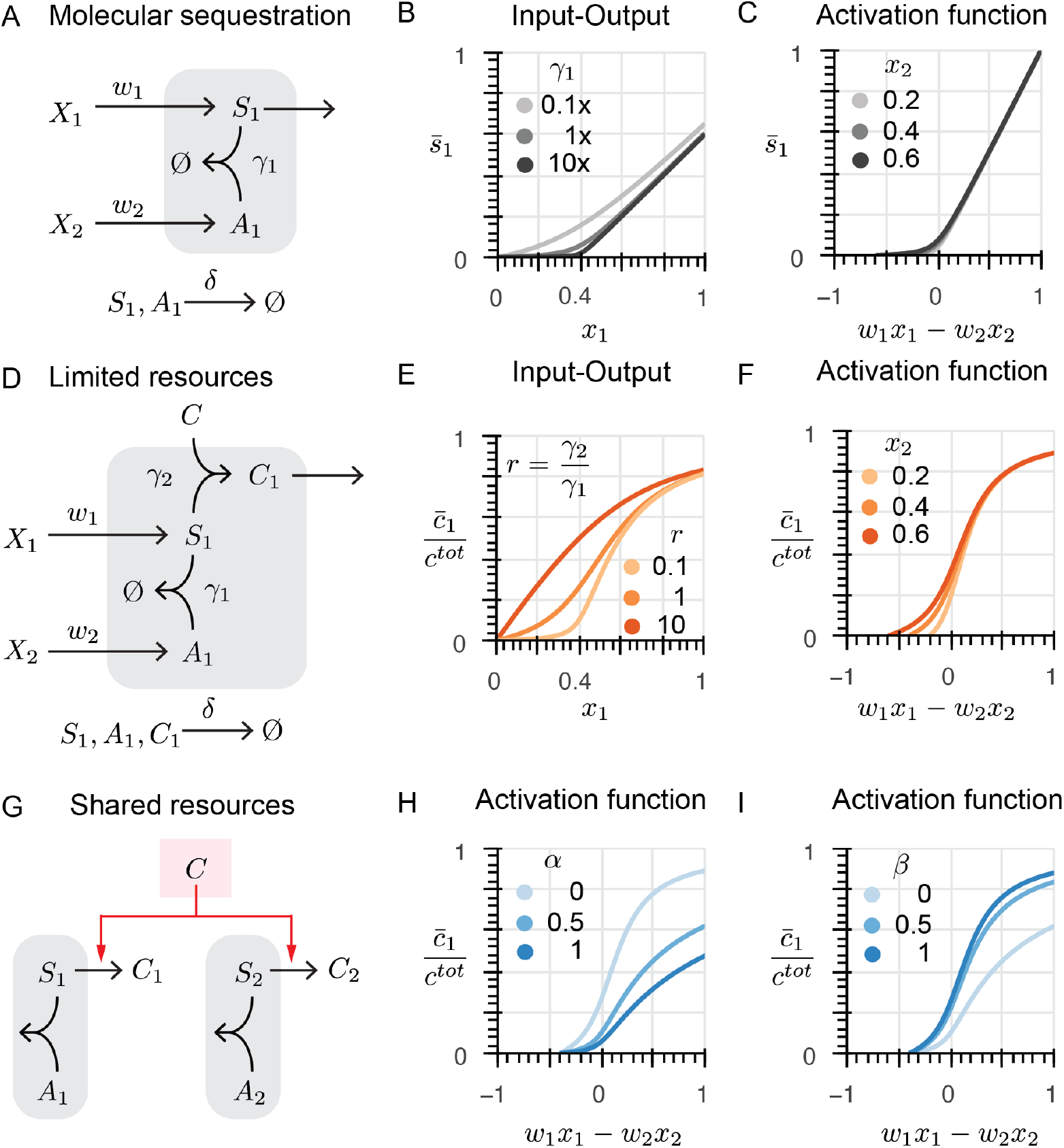
Molecular sequestration of a sigma factor (S_1_) and its anti-sigma (A_1_) recapitulates variants of ReLU activation function. **A**: Schematic representation of chemical reactions depicting molecular sequestration of S_1_ factor and A_1_. The inputs are concentrations of species X_1_ and X_2_ that generate S_1_ and A_1_, respectively. The output is steady-state concentration of S_1_ shown as 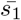. **B**: The effect of sequestration rate (γ_1_) on input-output relationship of the sequestration reaction. **C**: The relationship between 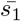 and the inputs x_1_ and x_2_ resembles a soft ReLU function. **D**: Schematic representation of chemical reactions depicting molecular sequestration of S_1_ and A_1_ as well as complex formation of S_1_ with a limited amount of core RNA polymerase (C). The inputs are concentrations of species X_1_ and X_2_ that generate S_1_ and A_1_, respectively. The output is steady-state normalized concentration of S-C dimer (C) shown as 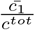. **E**: The ratio of RNApol-S complex formation rate to sequestration rate (referred to as competitive binding ratio r) influences the input-output relationship in the sequestration system. Lower r allows construction of a thresholding function. **F**: In fast sequestration regime and slow complex formation (r → 0 referred to as fast competitive binding regime), the output of the sequestration reactions in the presence of limited resources resembles an asymptotic saturated ReLU (ASReLU). **G**: Schematic representation of chemical reactions depicting molecular sequestration of S_1_ and A_1_ as well as complex formation of S_1_ with a limited amount of core RNA polymerase (C) in the presence of a competing sigma factor (S_2_). The inputs are concentrations of species X_1_ and X_2_ that generate S_1_ and A_1_, respectively. The output is steady-state normalized concentration of S-C dimer (C) shown as 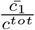. **H**: The effect of concentration of species α that generates the competing sigma factor S_2_ on the activation function of sequestration reaction. The addition of competitive binding reduces the amount of total C available, thus lowering the saturation level of the ASReLU function. **I**: The effect of concentration of species (β) that generates A_2_ that sequesters the competing S_2_ on the activation function of sequestration reaction. Higher sequestration of S_2_ results in a higher concentration of available C, thus increasing the saturation level of the ASReLU function.

Depending on the sequestration rate (*γ*_1_), the output of the system, 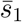, which stands for the steady-state amount of *S*_1_, follows the input (*w*_1_*x*_1_) in different patterns (**Fig. 2B**). To evaluate two regimes of output at the steady state, we define a dimensionless positive parameter 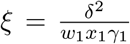. In the slow sequestration regime, when the affinity of *S*_1_ binding to *A*_1_ is small (*ξ* ≫ 1), the amount of 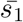 has a non-linear relationship with the input *x*_1_ (**Fig. 2B**). However, in the fast sequestration regime (when the binding affinity of *S*_1_ and *A*_1_ is large) when 0 *< ξ* ≪ 1, modeling the interaction between *S*_1_ and *A*_1_ (see section 1.1 in SI for mathematical derivation) leads to a thresholding function[31] (a function that generates positive outputs only when the input is larger than a threshold) corresponding to a linear relationship shown in equation (1).

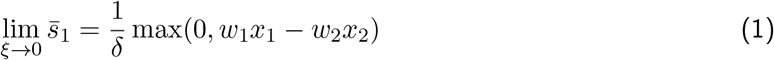

The assumption of fast sequestration of a sigma factor by its corresponding anti-sigma molecule is valid as previous studies on various anti-sigma molecules have found their rapid effect on transcriptional activity of their target sigma factors [36, 37, 38]. In addition, molecular controllers based on fast sigmaanti-sigma interaction have been constructed and tested [39, 40, 41], thereby providing evidence for fast sequestration assumption.

In equation (1), *x*_1_ and *x*_2_ represent the concentration of input species that generate *S*_1_ and *A*_1_ with production rates of *w*_1_ and *w*_2_, respectively. At steady-state and a fast sequestration regime, equation (1) converges asymptotically to a ReLU function. Therefore, we named the relationship between 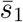 with the inputs *x*_1_ and *x*_2_ the Asymptotic ReLU (AReLU) function (**Fig. 2C**). Thus, the sequestration relationship between *S*_1_ and *A*_1_ resembles a simple perceptron with an AReLU activation function and weights of *w*_1_ and −*w*_2_ for inputs *x*_1_ and *x*_2_, respectively. The linear relationship regime between 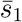 and *x*_1_ indicates that the steady-state available amount of *S*_1_ is simply the difference between the total steady-state amounts of *S*_1_ and *A*_1_, similar to other sequestration-based calculators demonstrated previously [42]. In other words, the outcome of the sequestration chemical reaction simply calculates the subtraction of total *S*_1_ and *A*_1_ and does not produce any product if *S*_1_ is lower than *A*_1_. However, in slow sequestration, the amount of 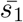 depends on the input *x*_1_ through a non-linear relationship similar to an activation function known as soft ReLU function [43] (**Fig. 2C**).

In natural systems, sigma factors either bind to their corresponding anti-sigma or the RNA polymerase (RNApol) core. In the latter case, the RNApol-sigma factor complex binds to a specific sigma promoter in the DNA sequence and drives the expression of the downstream genes. When free *S*_1_ is present, it binds to the RNApol to initiate transcription. Therefore, when designing sigma-based neural networks, since the amount of RNApol-sigma factor complex directly influences protein expression, the interaction between *S*_1_ and the RNApol core should be considered. Assuming that the sigma factor *S*_1_ binds to a limited amount of available RNApol core *C*, with total concentration denoted as *c*^*tot*^, at rate *γ*_2_, the total amount of RNA pol-sigma factor complex, denoted as *C*_1_, can be calculated by solving the system of ordinary differential equations (ODEs) representing the following chemical reactions which are illustrated in **Fig. 2D**:

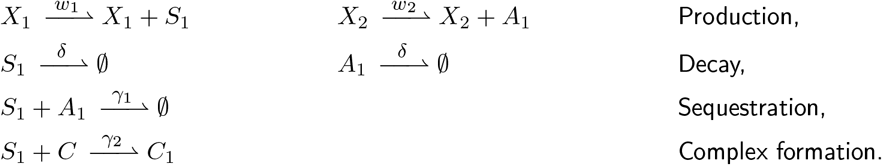

To analyze the input-output relationship of the chemical reactions that consider limited resources, we introduce a dimensionless variable 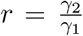 referred to as competitive binding ratio that represents the ratio of sigma factor-RNApol affinity to the sequestration rate. Solving the ODEs representing the above reactions (see section 1.2 in SI for derivation) reveals that the relationship between steady-state *C*_1_ denoted as *c*_1_ (and consequently its normalized value denoted as 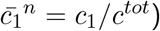 and *x*_1_ depends on the competitive binding ratio. When *r* ≫ 1 (slow competitive binding regime) the input-output relationship deviates from the ideal behavior (thresholding function) observed in **Fig. 2B**. However, when *r* ≪ 1 (fast competitive binding regime), 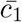 follows the thresholding function (**Fig. 2E**). Hence, when input *x*_1_ is lower than the threshold, there is no response, but when input is higher than the threshold, the response is non-zero and ultimately saturates at 1 (*c*^*tot*^).

Equation (2) describes the input-output function in the fast competitive binding regime (*r* ≪ 1).

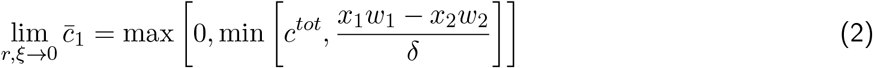

Further, equation (2) is closely similar to AReLU, equation (1), with the difference of having a limit on the output (*c*^*tot*^) which is introduced due to the limited resources. Thus, the effect of limited resources, in this case, total RNApol core, causes the deviation of 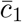 from a linear trend when *x*_1_ and *x*_2_ vary. Nevertheless, in the fast competitive binding regime even with a limited RNApol core, the activation function still performs as a subtraction calculator although it is saturated at *c*^*tot*^ (**Fig. 2F**). Equation (2) characterizes this non-linear relationship of 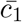 with inputs *x*_1_ and *x*_2_. We call this activation function asymptotic saturated ReLU (referred to as ASReLU hereafter).

Despite the ASReLU maintaining its function as a subtractor when the resources are limited, it is rarely the case for substrates to interact with their binding partners without other competing factors. For example, various sigma factors may compete with each other to bind to the available RNApol core molecules. Thus, when this competition is considered, the function of ASReLU may change. A simplified model consisting of two competing sigma factors (*S*_1_, *S*_2_) and their corresponding anti-sigma factors can be utilized to predict the function of ASReLU in the presence of competing sigma factors (**Fig. 2G**). For simplicity, here we assume that the competing sigma factors have identical affinities for their corresponding anti-sigma proteins as well as *C*. The effect of different kinetics will be investigated in the next section. The following chemical reactions represent the model:

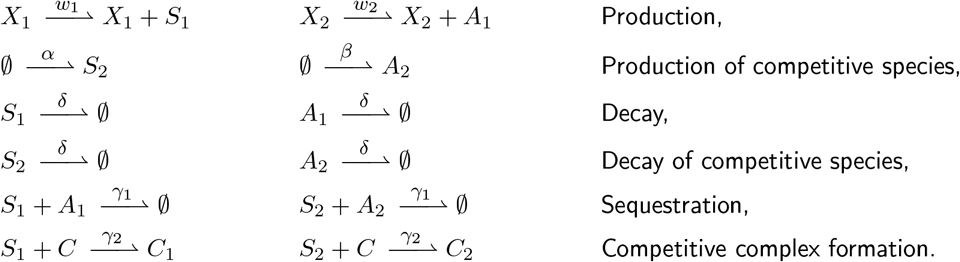

Solving at steady state, the set of ODEs modeling above equations yields equation (3) (for mathematical derivation, see SI section 1.3), which describes an expression of 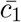 as a function of inputs (*x*_1_ and *x*_2_) and *c*^*tot*^. Although it is challenging to find a closed-form expression of 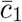 at the steady state, we can find an expression of 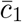 as a function of the inputs *x*_1_, *x*_2_, and 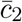 (unknown). This expression is useful to understand the effect of coupling between the two competing sigma factors on their binding to the RNApol. Equation (3) shows that the available RNApol is depleted by the other sigma factor reflected in the term 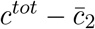. Additionally, the amount of depletion of available resources will depend on the production rates of both the competing sigma factor and its anti-sigma protein. Yet, in the fast sequestration regime (*ξ* → 0), and fast competitive binding (*r* → 0), the system converges to an ASReLU function with a lower saturation magnitude captured by 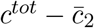.

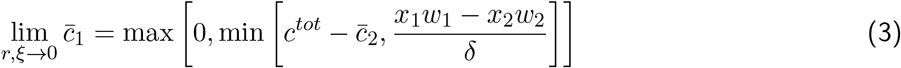

Since 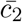 only influences the ASReLU saturation level, the performance of the ASReLU function in calculating the difference between *S*_1_ and *A*_1_ as a thresholding function is sustained under the effect of a competing sigma factor (**Fig. 2H**). However, the limit of output 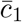 (the saturation level) further decreases as the amount of competing sigma factor *s*_2_ increases which corresponds with higher *α* (**Fig. 2H**). This trend reverses back to the non-competing ASReLU (equation (2)) when the amount of anti-sigma factor *A*_2_ increases with higher *β* since less *S*_2_ is available to compete with *S*_1_ (**Fig. 2I**).

Overall, molecular sequestration in a sigma factor-dependent translation system models variations of the ReLU function. In ideal conditions where the RNApol core is unlimited, this dependency is perfectly linear and reflects the difference between *S*_1_ and *A*_1_, enabling an AReLU function in a fast sequestration regime. However, in the presence of limited as well as shared resources, the trend between the sigma factor-RNApol complex and the differential value of *S*_1_ and *A*_1_ deviates from a linear trend and this deviation becomes more intense as the total amount of free *S*_2_ increases. This relationship in fast sequestration and competitive binding regimes is captured by the ASReLU function. However, since ASReLU still represents the difference of *S*_1_ and *A*_1_, we reasoned that the activation function can be used to create a perceptron, thus paving the way for constructing multi-layer neural networks for generating non-linear outputs.

### 2.2 Design and analysis of a sigma-based perceptron with ASReLU activation function and sharing limited resources

After confirming the output of the sigma-based perceptron with an ASReLU activation function, we seek to determine whether this system demonstrates typical characteristics of a single node in a neural network in the presence of a competing node. We test various input ranges of the perceptron and its competing counterparts to investigate the perceptron’s linear decision boundary as well as weight-tuning for manipulating its decision boundary. Thus, we model the binding of a sigma factor (*S*_1_) to RNApol in the presence of its anti-sigma (*A*_1_) as well as a competing sigma-anti-sigma pair (*S*_2_ and *A*_2_) and look at the steady-state total amount of RNApol bound to *S*1 (denoted as 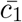) as the output of the node in response to a wide range of inputs (Figs. 3A and B).

**Figure 3:**
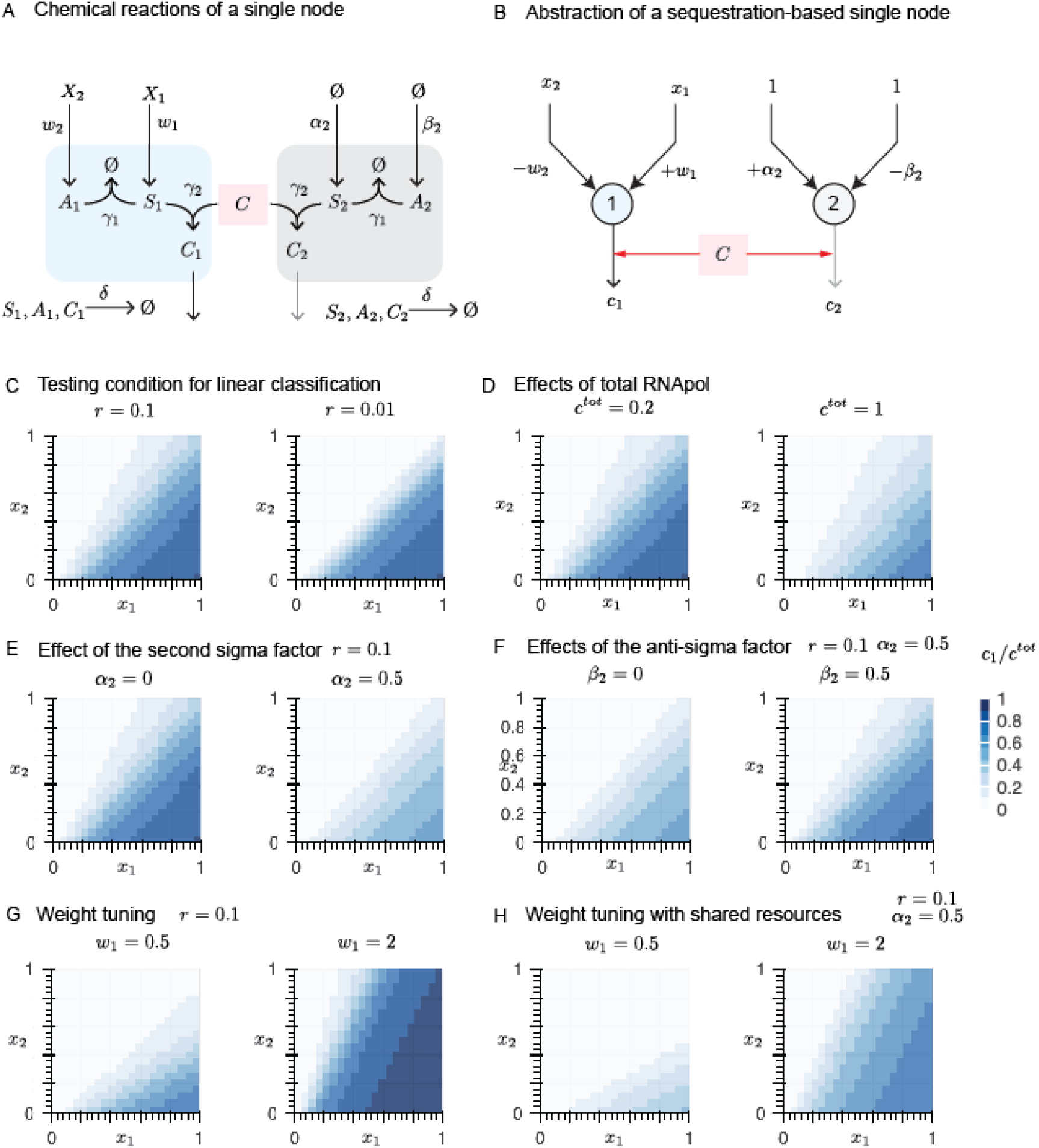
Biomolecular network with all or none response demonstrate characteristics of a perceptron. **A**: Chemical reactions representing a single perceptron in the presence of a competing node. See table 1 for the value of input weights. **B**: The equivalent representation of the chemical reactions in the context of artificial neural networks. **C**: Linear decision boundary of a single node in the absence of competition. Lower r demonstrates linear decision boundary while higher r generates linear decision boundaries with weights that vary as a function of input. **D**: Linear decision boundary of a single node with different amounts of total C in the absence of competition. A higher concentration of C allows a higher range of responses and lack of saturation observed with lower c^tot^. **E**: Linear decision boundary of a perceptron in the presence of a competing node. α_2_ represents the concentration of species that produces competing S_2_. A higher amount of S_2_ reduces the available concentration of C, thus suppressing the response of the decision boundary while leaving its pattern intact. **F**: The effect of concentration of A_2_ (controlled by β_2_) which sequesters competing S_2_ on the linear decision boundary of a perceptron in the presence of limited resources and a competing node. Higher sequestration yields a higher available concentration of C, thus increasing the amplitude of perceptron response. **G**: Linear decision boundary of a single node in the absence of a competing factor can be tuned by adjusting the weight of the input even when r = 0.1. **H**: Tuning the linear decision boundary of the perceptron in the presence of a competing node can still be realized by changing the input weight. The amplitude of the response, however, is suppressed due to the lower availability of C.

First, we investigate the output of the single node in isolation (without competing factors) to understand the effect of binding kinetics between *S*_1_ and *A*_1_ (*γ*_1_) as well as *S*_1_ and *C* (*γ*_2_) on the perceptron’s decision boundary. Our findings show that the competitive binding ratio plays a critical role in determining the response pattern, also known as the decision boundary, of the perceptron. Equation (4) below provides insight on how *r* and total resources *c*^*tot*^ affect the decision boundary (see SI section 1.2 for derivation):

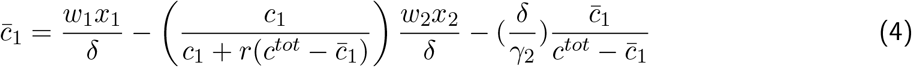

The last term in the equation (4) simply introduces a bias to the decision boundary which only influences the response amplitude while leaving the response pattern, dictated by weights *w*_1_ and *w*_2_, intact. Additionally, the coefficient of *x*_2_ depends on both variables *w*_2_ and 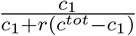. Therefore, the competitive binding ratio, depending on its magnitude, can change the slope of the decision boundary. When *r* → 0 (**Fig. 3C**, right), the coefficient of 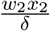 in equation (4) converges to 1. Hence, in this regime, the pattern of decision boundary remains linear across the input ranges although its bias changes depending on the inputs. However, as *r* becomes greater, the effect of limited resources on the decision boundary strengthens (**Fig. 3C**, left). In such a condition, the decision boundary still remains linear, but since *c*_1_ is a function of *x*_1_ and *x*_2_, the slope of the decision boundary varies across the range of the inputs as an inverse function of *c*^*tot*^ − *c*_1_. Intuitively, when *r* → 0, the sequestration rate *γ*_1_ is higher than *γ*_2_. Thus, the available amount of *S*_1_ becomes equal to *x*_1_*w*_1_ − *x*_2_*w*_2_ which corresponds to excess amount of *S*_1_ after binding to *A*_1_. In this case, regardless of the binding rate between *S*_1_ and the RNApol *C*, the output will be a simple linear function of excess *S*_1_. On the other hand, when the sequestration rate *γ*_1_ is slower than the binding rate between *S*_1_ and *C* (*γ*_2_), the dynamics of binding between *S*_1_ and *A*_1_ in conjunction with binding of *S*_1_ and *C* dictates non-linearity on the output of the sequestration and complex formation reactions (seen in **Fig. 2**F). The decision boundary of the perceptron, consequently, consists of linear decision boundaries with slopes that change as the inputs vary. Nevertheless, as long as *r* ≪ 1, the perceptron still functions similarly to the ideal condition (when *r* → 0) and generates a decision boundary which resembles the ideal linear condition and can be used for the construction of more complicated architectures.

Next, we seek to investigate how the total amount of RNApol, *C*, affects the perceptron function. Since *C* does not play a role in the dynamics of the ASReLU function, its variation reveals itself as a simple increase or decrease in perceptron response amplitude (**Fig. 3D**). This effect occurs because the normalized output of the ASReLU, 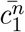, is completely independent of *C* and varies only by *x*_1_ and *x*_2_ (see equation (23) in the SI). Therefore, a single perceptron node in the absence of any competing sigma factors and in the presence of limited resources still acts as a linear classifier. However, the dynamic effects of anti-sigma-sigma binding on the system effectively change the weights of the perceptron decision boundary. Such changes, however, are minimal when *r* ≪ 1.

So far, we have focused on the decision boundary of the biological perceptron made by a sigma factor and its corresponding anti-sigma protein. Next, we consider whether the presence of another sigma factor (*S*_2_) and its anti-sigma protein (*A*_2_) changes the pattern of the perceptron decision boundary. Equation (5) provides an expression for the output of the perceptron as a function of inputs, *c*_1_, and *c*_2_. (See SI section 1.3 for derivation.)

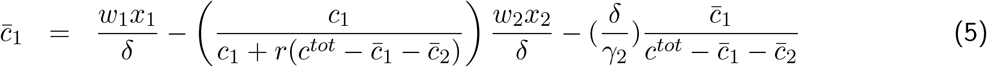

Equation (5) can be interpreted as an alternative form of equation (4) if *c*^*tot*^ in equation (4) is replaced with 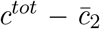. In other words, since the presence of a competing perceptron shrinks the amount of available resources, it amplifies the bias introduced to the perceptron output (by lowering the magnitude of the denominator in the last term in equation (5)) and strengthens the input-dependent slope variation in decision boundary of the main perceptron (by increasing the coefficient of 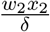 in equation (5)). Note that *c*_1_ in equation (5) is a function of inputs *x*_1_ and *x*_2_. Therefore, the magnitude of introduced bias and change in the slope of the decision boundary will be varied in different input regimes. However, if *r* ≪ 1, the effect of competition and resource sharing on tuning decision boundary becomes negligible as the coefficient of 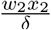 in equation (5) converges to 1.

Our simulations for a perceptron in a fast competitive binding regime over a range of different input concentrations verify our mathematical analysis by demonstrating that in the presence of another sigma factor (produced by input *α*_2_) competing to bind to the RNApol, the perceptron response is suppressed due to the bias introduced by the competing perceptron which lowers the amount of available RNApol (**Fig. 3E**). The response pattern, on the other hand, remains mainly intact due to the low competitive binding ratio.

We also investigate how anti-sigma *A*_2_ (produced by input *β*_2_) influences the response of the perceptron in a resource-sharing system. Binding of *A*_2_ to *S*_2_ disables *S*_2_ from binding to RNApol, thereby increasing the total amount of RNApol available for *S*_1_ to bind. Therefore, we expect that the introduction of *A*_2_ to the system suppresses the bias effect of *S*_2_ on the perceptron response pattern (last term in equation (5)). Indeed, our simulations show that an increase in production of *A*_2_ increases the perceptron output (**Fig. 3F**), thus confirming our hypothesis.

Our mathematical analysis of the perceptron output in the presence of a competing perceptron leads to an expression for 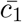 that is dependent on inputs and 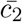 (equation (48) in the SI), indicating that kinetics of binding between *S*_2_ and *A*_2_ would reveal its effect on perceptron output by simply varying its response amplitude while leaving its response pattern intact as long as the perceptron functions in the fast competitive binding regime (see SI section 1.3 for mathematical derivation. See **Figs. S2 and S3** for simulations over a wide range of different *S*_2_ and *A*_2_ concentrations and different kinetics of *S*_2_-*A*_2_ binding, respectively.).

Lastly, as weight tuning is a fundamental characteristic of nodes in neural networks that defines their individual decision boundaries, we determine the feasibility of tuning the weights applied to our biological perceptron in the presence or absence of resource sharing. We first analyze an individual node with a limited amount of *C* and observe the change in the output pattern as *w*_1_ increased (**Fig. 3G**). Similar to perceptrons used in neural networks, adjusting the weight applied to the input results in a discernible change in the slope of the response pattern which is in agreement with our mathematical analysis (see 1.3 in SI). Aligned with our expectation, introducing a second node *S*_2_ that imposes resources sharing on the system preserves the weight-tuning characteristic of the perceptron and only affects its response amplitude by lowering the saturation limit (**Fig. 3H**). Taken together, our simulations elucidate the amplitude-modulation effect of a competing sigma factor on the perceptron output in the fast competitive binding regime and also demonstrate the weight-tuning ability of perceptron with or without resource sharing. Therefore, we concluded that by operating in a fast competitive binding regime, sequestration-based perceptrons can demonstrate linear decision boundaries despite sharing limited resources which in turn allows the construction of multi-layer perceptron networks.

### 2.3 Biomolecular neural networks generate non-linear classifiers in the presence of shared resources

While a single sigma-based perceptron generates a tunable decision boundary even with sharing limited resources (**Fig. 3**), most biologically relevant biocomputation processes such as competitive ligand binding and protein dimerization generate sophisticated non-linear responses that rely on the protein-protein interactions and the concentration of competing dimerizing proteins or ligands [18, 17]. Therefore, we aim to utilize the sigma-based perceptron as a basic building block of more intricate networks made of multiple perceptrons that are capable of generating non-linear outputs.

First, we investigate a two-node network where each node receives two inputs with unique weights (**Fig. 4A**). This simple network allows us to study the effect of sharing limited resources on the output of nodes in the same layer (see SI section 2.1 for mathematical representation of the network). In the absence of resource sharing, each node’s output is a linear combination of its inputs (**Fig. 3C**). Therefore, with given weights denoted in **Fig. 4A**, linear patterns for outputs of nodes 1 and 2 are observed (**Fig. 4B**). However, when the binding competition between sigma factors is taken into account, the effect of limited resources and competition on the network output is elucidated as the attenuation of outputs for each node in certain input regimes (**Fig. 4C**) resulting in a non-linear response pattern. Notably, the response of each node is maximally affected where the competing node is consuming most of the resources. For instance, in isolation, the output of node 2 (*c*_2_) has its highest expression level when [*x*_1_, *x*_2_] → 1 (**Fig. 4B**, bottom). Consequently, the response pattern of *c*_1_ is highly diminished in that region (**Fig. 4C**) due to the effect of resource sharing with node 2. In a similar fashion, the output pattern of *c*_2_ demonstrates its highest attenuation where *C*_1_ is expressed the most in isolation (*x*_1_ → 1), thus leaving a smaller amount of available RNApol (*C*) for production of *C*_2_. Hence, the concerted effect of resource sharing in a wide range of inputs imposes non-linearity in the output of nodes in the same layer due to the emergence of a dominant perceptron that consumes most of the resources. This effect can also be deduced from equation (5) knowing that both *c*_1_ and *c*_2_ are functions of inputs. Therefore, in certain input regimes where either *c*_1_ or *c*_2_ are strongly expressed, the effects of competitive resource sharing detailed in the previous section become strengthened.

**Figure 4:**
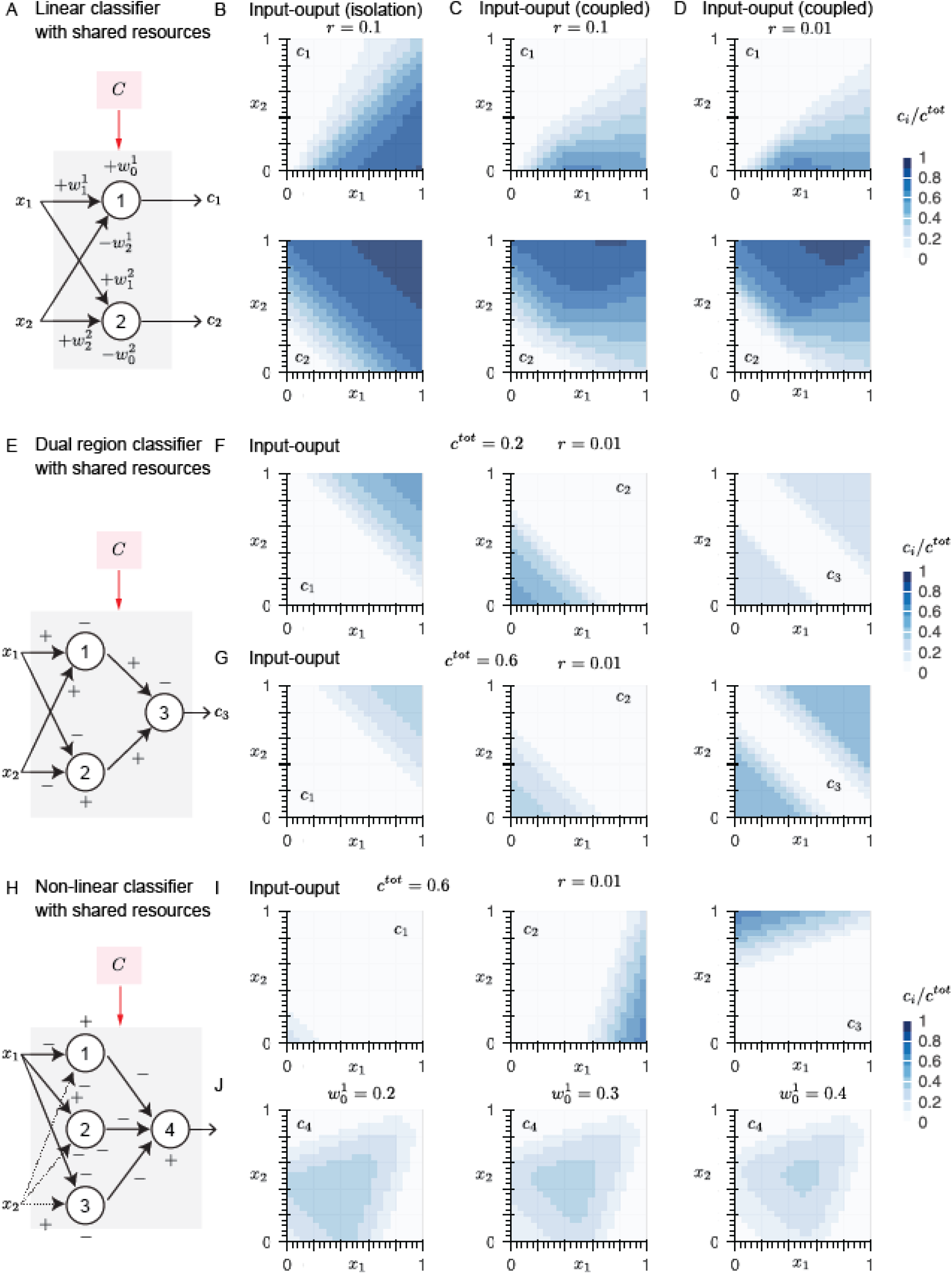
Biomolecular neural networks enable implementation of linear and non-linear classifiers: **A**: Schematic representation of a single-layer neural network that generates a linear classifier. Both perceptrons share the limited resources C. See table 1 for the value of input weights. **B**: Linear decision boundary of perceptron 1 (top) and 2 (bottom) in the absence of competitive binding. **C**: Decision boundary of perceptrons 1 (top) and 2 (bottom) when perceptrons compete to bind to the limited resources C. The competitive binding ratio r is 0.1. **D**: Decision boundary of perceptrons 1 (top) and 2 (bottom) when perceptrons compete to bind to the limited resources C. The competitive binding ratio r is 0.01. Competitive binding introduces non-linearity to the classifier by reducing the output amplitude in input regimes in which both perceptrons require resources. **E**: Schematic representation of the architecture of the dual region classifier multi-layer neural network with shared resources. + and − signs indicate the weights of inputs and whether they promote the expression of sigma or anti-sigma proteins. See table 1 for the value of input weights. **F**: Output maps of the perceptrons in the first layer (left and middle for c_1_ and c_2_, respectively) each representing a linear classifier with minimal interference. Right panel depicts network output (c_3_) which comprises the dual region classifier with c^tot^ = 0.2. **G**: Output maps of the perceptrons in the first layer (left and middle) and network output (right) which comprises the dual region classifier with c^tot^ = 0.6. The pattern of the classifier is invariant to c^tot^ while its amplitude varies with c^tot^. **H**: Schematic representation of the multi-layer neural network with a non-linear classifier output. + and − signs indicate the weights of inputs and whether they promote the expression of sigma or anti-sigma proteins. See table 1 for the value of input weights. **I**: The output of perceptrons in the first layer of the network. Perceptrons 1 (left), 2 (middle), and 3 (right) each recapitulate a linear classifier with zero to little interference in their linear decision boundaries. **J**: The output of the network depicted in **H** (c_4_) is a non-linear classifier. The classifier boundary can be tuned by adjusting the biases of nodes in the first layer. Shown are examples of different decision boundaries by changing the bias of perceptron 1 to 0.2 (left), 0.3 (middle), and 0.4 (right).

Interestingly, the network output is influenced differently in fast and slow competitive binding regimes. In a fast competitive binding regime, 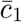 and 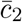 are linear functions of *x*_1_ and *x*_2_. Therefore, the outputs *c*_1_ and *c*_2_ display sharper deviations when both are dependent on inputs (**Fig. 4D**) due to the stronger effect of the bias term in equation (5) imposed by sharp reduction of resources (see **Fig. S4** for isolated perceptron responses in fast competitive binding regime.). Overall, we conclude that resource sharing can significantly induce non-linearity to the network output pattern and the highest impact of competitive resource sharing reveals itself where the response of isolated nodes overlap with each other, corresponding to input regimes with maximum resource sharing.

Knowing that overlapping responses can cause non-linearity in the network outputs, we next consider whether we can still implement non-linear networks with decision boundaries that are distorted minimally due to resource sharing. We reason that if the outputs of the same layer do not overlap with each other, the overall output of the network will follow the design principles dictated by the input weights and will not have undesired non-linearity. To test this hypothesis, we look at a simple network made of three nodes that reconstitutes a dual region classifier (band-stop filter) represented in **Fig. 4E** (see SI section 2.2 for mathematical representation of the network and **Fig. S5** for the ideal response of this network in the absence of sharing limited resources). In order to avoid non-linearity induced by resource sharing, we design the network such that the node outputs have minimal overlap with each other (**Fig. 4F**, left and middle panels). We expect that the lack of overlap between node outputs would prevent unwanted non-linearity in the network output (*c*_3_). Aligned with our expectation, our simulations demonstrate the network response consisting of two separate linear regions despite resource sharing (**Fig. 4F**, right).

We also confirmed that an increase in the amount of available resources (*C*) decreases the effect of competition for resources and does not alter the decision boundary of first layer nodes (**Fig. 4G**, left and middle) as well as overall network (**Fig. 4G**, right). Therefore, we conclude that linear responses can be engineered in a network, despite the presence of resource sharing, by tuning input weights such that the outputs of the same layer do not overlap.

Finally, we seek to create a biomolecular neural network that generates a non-linear classifier. Such non-linear outputs are key features of many biological processes and drivers of cell decision-making [44, 45, 46]. We design a simple network with specific input weights based on the interaction of multiple sigma factors and their corresponding anti-sigma molecules that generates a non-linear classifier resembling a band-pass filter (**Fig. 4H**, see SI section 2.3 for ODEs describing the network). As in the previous design, we want the outputs of the first layer to have minimal overlap with each other to avoid non-linearity induced by resource sharing. Our simulations depict that the first layer outputs, *c*_1_, *c*_2_, and *c*_3_ have minimal overlap with each other given the particular weights and biases (**Fig. 4I**). Consequently, our simulations demonstrate that network output (*c*_4_) creates the non-linear classifier (**Fig. 4J**) that is aligned with our ideal design expectations (**Fig. S6**). Consistent with our findings in the previous section, changing the *c*^*tot*^ does not change the non-linear pattern of the decision boundary but tunes its amplitude (**Fig. S7**).

We also look at the effect of input bias on the first layer of the output (**Fig. 4J**). Indeed, in accordance with the ideal design (**Fig. S6**), increasing the bias of node 1 amplifies its output (*c*_1_) which subsequently further sequesters the amount of available *S*_4_, thus suppressing the network response in its corresponding input regime ([*x*_1_, *x*_2_] → 0) without significant deviation from design expectations (**Fig. S6**). Therefore, we showed that even in the presence of resource sharing, we can develop non-linear biological neural networks to realize dual region and non-linear classifiers. We also demonstrated that we can modulate the network response by tuning the model parameters that are independent of limited resources.

## 3 Discussion

Biological signal processing units that reconstitute molecular linear and non-linear classifiers are powerful tools that enable cellular decision-making, precise cell programming, highly discriminatory input processing, and ultra-sensitive molecular biosensors for applications such CAR T-cell engineering or *in vitro* diagnostics. A biological perceptron ideally allows linear classification while a combination of biological perceptrons can create biomolecular neural networks that compute non-linear classification.

Among different approaches that are used to construct biochemical neural networks and classifiers, sequestration-based networks are of significant interest because of their simplicity and compatibility with *in vivo* systems [31, 24]. However, the effect of physiological constraints such as a limitation on resources, as well as competitive binding between elements of molecular classifier networks and specifically sequestration-based networks on the classification function have remained unexplored.

We mathematically modeled sequestration-based biochemical neural networks and investigated how sharing limited resources, a ubiquitous feature of physiological systems, influences the function of the neural network decision boundary. We chose a network of sigma transcription factors, their corresponding anti-sigma proteins, and core RNA polymerase as our model system. Our modelings demonstrated that a single perceptron, the basic building block of neural networks, with a ReLU-like activation function is recapitulated by modeling the interaction of one sigma factor with its anti-sigma molecule. We further showed that modifying the model to include a limited amount of core RNApol in the system changes the activation function of the sigma-based perceptron to an asymptotic saturated ReLU. Drawing inspiration from natural systems where multiple sigma factors compete to bind to a limited pool of RNApol, we altered the model to include another sigma factor and found that the decision boundary of the sigma-based perceptron remains the same although its output is suppressed.

While conditions of biochemical *in vitro* reactions are primarily controlled, in living organisms, endogenous factors can cause perturbations and deviations from the ideal design. For instance, in bacteria, although a certain sigma factor might be designed to control gene expression, multiple other sigma factors compete with the engineered sigma factor to bind to a limited amount of RNApol. To include competitive binding to shared resources, we increased the number of perceptrons, controlled by the same inputs, to two. We found that in specific input regimes where the outputs of the perceptrons interfere with each other, one dominant perceptron emerges and consumes most of the resources whereas the decision boundary of the non-dominant perceptron significantly deviates from its ideal design.

Given that engineering any kind of non-linear response by neural networks requires multi-layer perceptrons, we investigated conditions where despite resource sharing, non-linear decision boundaries could be designed. Knowing that interference of outputs of perceptrons in the same layer raises deviations of the linear decision boundary, we engineered particular multi-layer perceptrons in which non-linear decision boundaries were successfully demonstrated. We showed that despite sharing limited resources, dual region and non-linear classifiers resembling band-stop and band-pass filters can be implemented in sigma-based neural networks using different architectures of 2-layer neural networks with minimal deviation from ideal design.

An advantage of sigma-based sequestration-based neural networks over DNA-based neural networks is that they originate from endogenous proteins in bacteria. Therefore, sigma-based neural networks can be implemented both in bacterial systems as well as bacterial lysate-based cell-free expression systems, thus expanding their applications as living computers as well as *in vitro* biosensors and synthetic cell-based biocomputers[47, 48]. To indicate the possible implementation of neural networks demonstrated here (**Fig. 4**), we present schematics of required DNA sequences for each network in **Figs. S8** and **S9**. Additionally, unlike neuromorphic systems made of inducible genetic circuits, in sequestration-based networks, both positive and negative weights can be engineered, thereby making them more flexible and applicable for generating non-linear response patterns. Since outputs of sigma-based perceptrons are transcriptionally active sigma-RNApol complexes, they can be programmed to drive the expression of other sigma factors. Relying on this characteristic of sigma-based perceptrons, multi-layered perceptrons that are capable of recapitulating sophisticated non-linear response patterns can be designed readily. However, as shown in this work, for the neural network to function as designed and avoid perturbations caused by sharing limited resources among perceptrons, the outputs of perceptrons in the same layer must have minimal interference.

While we demonstrated designs of linear and non-linear classifiers in this work with predetermined weights that generate desirable decision boundaries, we note that sigma-based neural networks are not capable of learning through common algorithms like backpropagation. *i.e*., the input weights that directly determine the decision boundary are chosen by the designer. However, utilizing the mathematical analysis presented here, one can implement optimization approaches such as particle swarm optimization or other heuristic algorithms to find the appropriate weights for the generation of the desired decision boundary *in silico* prior to testing them *in vivo* or *in vitro*. Such optimization algorithms can also be implemented to expand the function of BNNs to regimes where non-linear effects of resource constraints appear. Therefore, engineering biologically constrained BNNs to recognize any arbitrary input pattern using optimization methods is an avenue worth exploring in the future.

Recently, it was shown that many cancer cell types can be recognized with higher precision if a combination of two antigens is used to identify them instead of traditionally using one biomarker[49]. Implementing non-linear output patterns with sequestration-based neural networks could increase the recognition ability of engineered cellular systems like CAR T-cells by equipping them with information-processing neural networks that generate desired outputs only in designed antigen concentration regimes. Similarly, by coupling inputs to the expression of sigma factors and their anti-sigma molecules, more sensitive, precise, and versatile *in vitro* biosensors for the detection of pathogens, substances, and biomarkers can be constructed.

With the nanofabrication technology in the semiconductor industry approaching its physical limits of manufacturing smaller and smaller elements [50], alternative computational devices with biological components are gaining increasing interest. However, current biocomputational systems are only in their infancy. Although biocomputation in living systems was initially shown more than two decades ago using genetic circuits, the limited range of computational tasks that genetic circuits can perform as well as the digital nature of their input-outputs makes their application limited. With the recent booming advance of AI, biochemical approaches that recapitulate biological neural networks holding the potential to perform intricate computational tasks have gained attention [3, 51]. Our study provides a general framework for designing biological perceptron or linear classifiers using existing biomolecular tools in the presence of resource constraints that are ubiquitous in physiological conditions. This framework, thus, is the first step towards designing sophisticated biomolecular neural networks that equip engineered cells with high-level computational and decision-making abilities.

In addition to transformative applications of sigma-based neural networks used in a forward-engineering manner in both cellular and cell-free systems, the fact that sigma and anti-sigma molecules can construct complicated computational modules elucidates the hidden capabilities of these rather simple transcriptional regulation molecules in cells. It was revealed by Park *et al*. that sigma factors share the resources in a pulsatile manner [20]. However, here we demonstrated that sharing limited resources influences sigma-based processes beyond time sharing. Bacterial cells have many sigma factors, some of which are activated only when their inputs meet certain conditions. However, how bacteria utilize the principles of molecular sequestration as well as sharing limited resources to respond differently to input combinations awaits future studies. Additionally, if bacteria are able to process certain input patterns using their endogenous sigma-based neural networks, the nature of these patterns and their role in guiding bacteria to make particular decisions remain unclear. In conclusion, our investigation demonstrates the effects of resource-sharing on sigma-based sequestration-based neural networks with up to three sigma factors and provides an outline for designing sigma-based non-linear neural networks in bacterial systems.

## 4 Acknowledgement

We thank Jean-Baptiste Lugagne, Noah Olsman, and Elisa Franco for helpful discussions. We thank Cold Spring Harbor Labs (CSHL) and instructors and TAs of CSHL synthetic biology course during summer 2022. Work in the A.P.L.’s laboratory has been supported by the National Science Foundation (EF1935265). H.M. thanks support from the University of Michigan Rackham Predoctoral Fellowship. I.G. acknowledges assistance from Boehringer Ingelheim Fonds. S.R.C. acknowledges funding from the Mayo Foundation for Medical Education and Research and Mayo Clinic’s Department of Molecular Medicine Small Grant. The 2022 CSHL Synthetic Biology course was funded by the National Science Foundation (MCB 2207222), with additional support from the Helmsley Charitable Trust and the Howard Hughes Medical Institute.

