## Supplemental information for "Engineering sequestration-based biomolecular classifiers with shared resources"

April 15, 2024

### 1 Steady state analysis of sequestration-based perceptrons

The following sections describe the derivation of ODES describing sigma-antisigma molecular sequestration and the construction of a perceptron based on their behavior. Where possible, analytical expressions are derived. All ODEs were modeled and the steady-state solutions were solved by a custom Python code using the numpy package. The figures were plotted using Bokeh and Matplotlib packages.

#### 1.1 Steady state of molecular sequestration in isolation

| | | |
|---|---|---|
| $X_1 \xrightarrow{w_1} X_1 + S_1$ | $X_2 \xrightarrow{w_2} X_2 + A_1$ | Production |
| $S_1 \xrightarrow{\delta} \emptyset$ | $A_1 \xrightarrow{\delta} \emptyset$ | Decay |
| $S_1 + A_1 \xrightarrow{\gamma_1} \emptyset$ | | Sequestration. |

Using the law of mass action, the above reactions are mathematically modeled into the following Ordinary Differential Equations (ODEs):

$$\dot{s}_1 = \alpha_1 - \delta s_1 - \gamma_1 s_1 a_1, \quad (1)$$

$$\dot{a}_1 = \beta_1 - \delta a_1 - \gamma_1 s_1 a_1, \quad (2)$$

where $\alpha_1 = w_1 x_1$, and $\beta_1 = x_2 w_2$.

At the steady state, by making equations (1), and (2) equal to zero, we can find the following equality:

$$\bar{a}_1 = \frac{\alpha_1 - \delta \bar{s}_1}{\gamma_1 \bar{s}_1} = \frac{\beta_1}{\gamma_1 \bar{s}_1 + \delta} \quad (3)$$

This leads to a second order polynomial, $P_1(\bar{s}_1) = \bar{s}_1^2 + A_1 \bar{s}_1 + B_1 = 0$, where $A_1 = \frac{\beta_1 - \alpha_1}{\delta} + \frac{\delta}{\gamma_1}$, and $B_1 = -\frac{\alpha_1}{\gamma_1}$. The polynomial $P_1$ admits a single positive real solution because $B_1$ is negative. This results in

$$\bar{s}_1 = \frac{1}{2}\left(-A_1 + \sqrt{A_1^2 - 4B_1}\right). \quad (4)$$

We can rewrite the above equation as

$$\bar{s}_1 = \left\{\frac{1}{2}\left((1 - \frac{\beta_1}{\alpha_1} - \xi) + \sqrt{(1 - \frac{\beta_1}{\alpha_1} - \xi)^2 + 4\xi}\right)\right\}\frac{\alpha_1}{\delta}, \quad (5)$$

where $\xi = \frac{\delta^2}{\alpha_1 \gamma_1}$. Further, we can approximate the steady state of $\bar{s}_1$ in the fast sequestration regime $\xi \to 0$ ($\gamma_1 >> \frac{\delta^2}{\alpha_1}$). This results in

$$\lim_{\xi \to 0} \bar{s}_1 = \max\left(0, 1 - \frac{\beta_1}{\alpha_1}\right)\frac{\alpha_1}{\delta} = \max\left(0, \frac{\alpha_1 - \beta_1}{\delta}\right) \quad (6)$$

#### 1.2 Steady state of molecular sequestration with limited resources

Here, we describe the interaction of sigma factor, anti-sigma factor and the RNA polymerize. We report all the chemical reactions of the sigma factor network.

| | | |
|---|---|---|
| $X_1 \xrightarrow{w_1} X_1 + S_1$ | $X_2 \xrightarrow{w_2} X_2 + A_1$ | Production |
| $S_1 \xrightarrow{\delta} \emptyset$ | $A_1 \xrightarrow{\delta} \emptyset$ | Decay |
| $S_1 + A_1 \xrightarrow{\gamma_1} \emptyset$ | $S_1 + C \xrightarrow{\gamma_2} C_1,$ | Sequestration/Complex. |

where $\alpha_1 = w_1 x_2$, and $\beta_2 = w_2 x_2$.

Using the law of mass action, the above reactions are mathematically modeled into the following Ordinary Differential Equations (ODEs):

$$\begin{aligned} \dot{s}_1 &= \alpha_1 - \delta s_1 - \gamma_1 s_1 a_1 - \gamma_2 s_1 c & (7) \\ \dot{a}_1 &= \beta_1 - \delta a_1 - \gamma_1 s_1 a_1 & (8) \\ \dot{c}_1 &= \gamma_2 s_1 c - \delta c_1. & (9) \end{aligned}$$

By making equations (7)-(9) equal to zero, we can find $\bar{c}_1$ at the steady state. First, we begin with $\dot{s}_1 = \dot{a}_1 = 0$ and find $\bar{a}_1$ as a function of $\bar{s}_1$:

$$\bar{a}_1 = \frac{\alpha_1 - \delta \bar{s}_1 - \gamma_2 \bar{s}_1 \bar{c}}{\gamma_1 \bar{s}_1} = \frac{\beta_1}{\gamma_1 \bar{s}_1 + \delta}. \tag{10}$$

This leads to the following expression,

$$\bar{s}_1 = \frac{\alpha_1}{\delta} - \left( \frac{\bar{s}_1}{\bar{s}_1 + \frac{\delta}{\gamma_1}} \right) \frac{\beta_1}{\delta} - \left( \frac{\bar{s}_1}{\bar{s}_1 + \frac{\delta}{\gamma_2}} \right) c^{tot}. \tag{11}$$

We can solve for $\bar{s}_1$, a third order polynomial $P_2(\bar{s}_1) = \bar{s}_1^3 + A\bar{s}_1^2 + B\bar{s}_1 + C = 0$ with coefficients $A = -\frac{\alpha_1}{\delta} + \frac{\beta_1}{\delta} + c^{tot} + \frac{\delta}{\gamma_1} + \frac{\delta}{\gamma_2}$, $B = -\frac{\alpha_1}{\gamma_1} - \frac{\alpha_1}{\gamma_2} + \frac{\beta_1}{\gamma_2} + \frac{\delta}{\gamma_1} c^{tot} + \frac{\delta^2}{\gamma_1 \gamma_2}$ and $C = -\frac{\delta \alpha_1}{\gamma_1 \gamma_2}$. Further, we can find an approximate solution of $\bar{s}_1$ at the steady state for $\gamma_2, \gamma_1 \to \infty$ (fast sequestration and complex formation rate). This results in

$$\lim_{\gamma_2, \gamma_1 \to \infty} \bar{s}_1 = \max \left[ 0, \frac{\alpha_1 - \beta_1 - \delta c^{tot}}{\delta} \right].$$

However, the analysis to understand the input-output behavior for $\bar{c}_1$ using the above expression is not simple. Instead, we will find an expression for $\bar{c}_1$. To do this, we use equation (9) at the steady state and find

$$\bar{c}_1 = \frac{\gamma_2}{\delta} \bar{s}_1 \bar{c}. \tag{12}$$

Then, we use mass conservation of c ($\bar{c} + \bar{c}_1 = c^{tot}$), and express $\bar{c}$ and $\bar{c}_1$ as a function of $\bar{s}_1$:

$$\bar{c} = \frac{1}{1 + \frac{\gamma_2}{\delta} \bar{s}_1} c^{tot} \tag{13}$$

$$\bar{c}_1 = \frac{\bar{s}_1}{\frac{\delta}{\gamma_2} + \bar{s}_1} c^{tot} \tag{14}$$

Further, we can use equation (14) to solve for $\bar{s}_1$, which leads to

$$\bar{s}_1 = (\frac{\delta}{\gamma_2}) \frac{\bar{c}_1}{c^{tot} - \bar{c}_1}. \tag{15}$$

Then, we can use equation (11) to find an expression for $\bar{c}_1$ by replacing $\bar{s}_1$ as a function of $\bar{c}_1$. This leads to

$$\bar{c}_1 = \frac{\alpha_1}{\delta} - \left( \frac{c_1}{c_1 + \frac{\gamma_2}{\gamma_1}(c^{tot} - c_1)} \right) \frac{\beta_1}{\delta} - (\frac{\delta}{\gamma_2}) \frac{\bar{c}_1}{c^{tot} - \bar{c}_1} \tag{16}$$

For the next steps, we normalize $\bar{c}_1$ by $c^{tot}$. Then, we define new variables $\alpha = \alpha_1/\delta/c^{tot}$, $\beta = \beta_1/\delta/c^{tot}$, $\xi = \delta/\gamma_2/c^{tot}$, $r = \gamma_2/\gamma_1$, and $\bar{c}_1^n = \bar{c}_1/c^{tot}$. As a result, we find the following normalized expression:

$$\bar{c}_1^n \;=\; \alpha - \left(\frac{\bar{c}_1^n}{\bar{c}_1^n + r(1-\bar{c}_1^n)}\right)\beta - \xi\frac{\bar{c}_1^n}{1-\bar{c}_1^n}. \tag{17}$$

This leads to a third order polynomial, $Q_2(\bar{c}_1^n) = A(\bar{c}_1^n)^3 + B(\bar{c}_1^n)^2 + C\bar{c}_1^n + D = 0$, where

$$A \;=\; 1 - r \tag{18}$$

$$B \;=\; (\beta - \alpha - \xi - 1) + (\alpha + \xi + 2)r \tag{19}$$

$$C \;=\; (\alpha - \beta) - (\xi + 1 + 2\alpha)r \tag{20}$$

$$D \;=\; \alpha r \tag{21}$$

We can rewrite the polynomial as

$$\bar{c}_1^n(\bar{c}_1^n - \alpha + \beta)(\bar{c}_1^n - 1) = h_1\xi + h_2 r + h_3\xi r, \tag{22}$$

where $h_1 = (\bar{c}_1^n)^2$, $h_2 = (\bar{c}_1^n - 1)^2(\bar{c}_1^n - \alpha)$, and $h_3 = (\bar{c}_1^n - 1)\bar{c}_1^n$

Depending on the value of $\alpha - \beta$, equation (22) can have different roots given that $\bar{c}_1$ can take values between $0$ and $1$. Fig. S1 indicates that the equation (22) has only one positive real root in the $[0, 1]$ interval regardless of the value of $\alpha - \beta$ which confirms the system converges to one steady-state solution. The roots of the left-side polynomial are $\bar{c}_1^1 = 0$, $\bar{c}_1^2 = \alpha - \beta$, and $\bar{c}_1^3 = 1$. Next, we focus on the regime of $r \to 0$ and how $O(r)$ affects the roots $\bar{c}_1^1 = 0$, and $\bar{c}_1^3 = 1$. When $\bar{c}_1$ gets close to 0 from the right side ($\bar{c}_1 \to 0^+$), $O(r)$ becomes negative and pushes the root to the left. This results in $\bar{c}_1^1 < 0$. Now, we focus when $\bar{c}$ gets close to 1 from the left side ($\bar{c}_1 \to 1^-$), $O(r)$ becomes small $\bar{c}_1^3$ and only $\delta\bar{c}_1$ will affect the solution and shift it to the right. This results in $\bar{c}_1^3 > 1$. Now, we can analyze $\bar{c}_1^2$ in three different regimes: negative ($\bar{c}_1^2 < 0$), larger than 1 ($\bar{c}_1^2 > 1$), and in the range of 0 to 1. When $\bar{C}_1^2$ is negative, $\bar{c}_1^1$ becomes positive ($\bar{c}_1^1 > 0$). When $\bar{C}_1^2$ is larger than 1, $\bar{c}_1^3$ becomes smaller than 1 ($\bar{c}_1^3 < 1$). Finally, when $\bar{c}_1^2$ is in between $0$ and $1$, $\bar{c}_1^2$ can be shifted to either left or right from $\alpha - \beta + \delta$, and this effect will depend on the parameters of the network. In summary, when $r, \xi \to 0$, and the solution is bounded between $0$ and $1$, we can approximate the only admissible solution as

$$\lim_{r,\xi\to 0} \bar{c}_1^n = \max[0, \min[1, \alpha - \beta]] \tag{23}$$

We can rewrite it as

$$\lim_{r,\xi\to 0} \bar{c}_1 = \max\left[0, \min\left[c^{tot}, c^{tot}\frac{\alpha_1 - \beta_1}{\delta}\right]\right] \tag{24}$$

#### 1.3 Steady state analysis of two sigma/anti-sigma factors competing for RNA polymerase

In this case, the following chemical reactions describe the system:

$$\begin{aligned}
&\emptyset \xrightarrow{\alpha_1} S_1 && S_1 \xrightarrow{\delta} \emptyset \\
&\emptyset \xrightarrow{\beta_1} A_1 && A_1 \xrightarrow{\delta} \emptyset \\
&S_1 + A_1 \xrightarrow{\gamma_1} \emptyset && S_1 + C \xrightarrow{\gamma_2} C_1 \\
&\emptyset \xrightarrow{\alpha_2} S_2 && S_2 \xrightarrow{\delta} \emptyset \\
&\emptyset \xrightarrow{\beta_2} A_2 && A_2 \xrightarrow{\delta} \emptyset \\
&S_2 + A_2 \xrightarrow{\gamma_1} \emptyset && S_2 + C \xrightarrow{\gamma_2} C_2
\end{aligned}$$

Using the law of mass action, we can derive the ODEs:

$$\dot{s}_1 = \alpha_1 - \delta s_1 - \gamma_1 s_1 a_1 - \gamma_2 s_1 c \tag{25}$$

$$\dot{a}_1 = \beta_1 - \delta a_1 - \gamma_1 s_1 a_1 \tag{26}$$

$$\dot{s}_2 = \alpha_2 - \delta s_2 - \gamma_1 s_2 a_2 - \gamma_2 s_2 c \tag{27}$$

$$\dot{a}_2 = \beta_2 - \delta a_2 - \gamma_1 s_2 a_2 \tag{28}$$

$$\dot{c}_1 = \gamma_2 s_1 c - \delta c_1 \tag{29}$$

$$\dot{c}_2 = \gamma_2 s_2 c - \delta c_2. \tag{30}$$

The total amount of $c$ is constant as a result we can write down the mass conservation of $c$ as $c_1 + c_2 + c = c^{tot}$.

At the steady state, we can find the following expressions:

$$\bar{a}_1 = \frac{\alpha_1 - \delta \bar{s}_1 - \gamma_2 \bar{s}_1 \bar{c}}{\gamma_1 \bar{s}_1} = \frac{\beta_1}{\gamma_1 \bar{s}_1 + \delta} \tag{31}$$

$$\bar{a}_2 = \frac{\alpha_2 - \delta \bar{s}_2 - \gamma_2 \bar{s}_2 \bar{c}}{\gamma_1 \bar{s}_2} = \frac{\beta_2}{\gamma_1 \bar{s}_2 + \delta} \tag{32}$$

$$\bar{c}_1 = \frac{\gamma_2}{\delta} \bar{s}_1 \bar{c} \tag{33}$$

$$\bar{c}_2 = \frac{\gamma_2}{\delta} \bar{s}_2 \bar{c} \tag{34}$$

Mass conservation allows us to solve for $\bar{c}$ and result in

$$\bar{c} = \frac{1}{1 + \frac{\gamma_2}{\delta} s_1 + \frac{\gamma_2}{\delta} s_2} c^{tot}. \tag{35}$$

Then, we can find $\bar{c}_1$ and $\bar{c}_2$ a function of $\bar{s}_1$ and $\bar{s}_2$,

$$\bar{c}_1 = \frac{s_1}{\frac{\delta}{\gamma_2} + s_1 + s_2} c^{tot} \tag{36}$$

$$\bar{c}_2 = \frac{s_2}{\frac{\delta}{\gamma_2} + s_1 + s_2} c^{tot}. \tag{37}$$

We use these two expressions to solve for $\bar{s}_1$ and $\bar{s}_2$ as a function of $\bar{c}_1$ and $\bar{c}_2$. As a result, we find

$$\bar{s}_1 = \frac{\delta}{\gamma_2} \left( \frac{\bar{c}_1}{c^{tot} - \bar{c}_1 - \bar{c}_2} \right) \tag{38}$$

$$\bar{s}_2 = \frac{\delta}{\gamma_2} \left( \frac{\bar{c}_2}{c^{tot} - \bar{c}_1 - \bar{c}_2} \right) \tag{39}$$

Next, we can plug in the below expression, From the steady-state expression for $\bar{s}_1$,

$$\bar{s}_1 = \frac{\alpha_1}{\delta} - \left(\frac{\bar{s}_1}{\bar{s}_1 + \frac{\delta}{\gamma_1}}\right)\frac{\beta_1}{\delta} - \left(\frac{\bar{s}_1}{\bar{s}_1 + \frac{\delta}{\gamma_2}}\right)c^{tot}. \tag{40}$$

This results in the following equality as a function of $\bar{c}_1$ and $\bar{c}_2$:

$$\bar{c}_1 = \frac{\alpha_1}{\delta} - \left(\frac{c_1}{c_1 + \frac{\gamma_2}{\gamma_1}(c^{tot} - \bar{c}_1 - \bar{c}_2)}\right)\frac{\beta_1}{\delta} - (\frac{\delta}{\gamma_2})\frac{\bar{c}_1}{c^{tot} - \bar{c}_1 - \bar{c}_2} \tag{41}$$

Following similar steps as previous section, we define $\alpha = \alpha_1/\delta/(c^{tot} - \bar{c}_2)$, $\beta = \beta_1/\delta/(c^{tot} - \bar{c}_2)$, $\xi = \delta/\gamma_2/(c^{tot} - \bar{c}_2)$, $r = \gamma_2/\gamma_1$, and $c_1^n = c_1/(c^{tot} - \bar{c}_2)$ (normalization). This leads to a third order polynomial, $Q_3(\bar{c}_1^n) = A(\bar{c}_1^n)^3 + B(\bar{c}_1^n)^2 + C\bar{c}_1^n + D = 0$, where

$$A = 1 - r \tag{42}$$
$$B = (\beta - \alpha - \xi - 1) + (\alpha + \xi + 2)r \tag{43}$$
$$C = (\alpha - \beta) - (\xi + 1 + 2\alpha)r \tag{44}$$
$$D = \alpha r \tag{45}$$

We can rewrite the polynomial as

$$\bar{c}_1^n(\bar{c}_1^n - \alpha + \beta)(\bar{c}_1^n - 1) = h_1\xi + h_2 r + h_3\xi r, \tag{46}$$

where $h_1 = (\bar{c}_1^n)^2$, $h_2 = (\bar{c}_1^n - 1)^2(\bar{c}_1^n - \alpha)$, and $h_3 = (\bar{c}_1^n - 1)\bar{c}^n$. In the fast sequestration and complex formation regime, we can approximate the steady state as

$$\lim_{r,\delta\to 0} \bar{c}_1^n = \max[0, \min[1, \alpha - \beta]] \tag{47}$$

Then, we can rewrite it as

$$\lim_{r,\delta\to 0} \bar{c}_1 = \max\left[0, \min\left[c^{tot} - \bar{c}_2, \frac{\alpha_1 - \beta_1}{\delta}\right]\right] \tag{48}$$

Similarly, following a similar analysis, we can find

$$\lim_{r,\delta\to 0} \bar{c}_2 = \max\left[0, \min\left[c^{tot} - \bar{c}_1, \frac{\alpha_2 - \beta_2}{\delta}\right]\right] \tag{49}$$

### 2 Steady-state analysis of sequestration-based biomolecular neural networks

The following subsections derive the ODEs describing the different biomolecular neural networks presented in **Fig. 4**. All ODEs were modeled and the steady-state solutions were solved by a custom Python code using the numpy package. The figures were plotted using Bokeh and Matplotlib packages.

#### 2.1 Two sigma-based networks that process the same inputs

We begin with the chemical reaction of the first node::

$$X_1 \xrightarrow{w_1^1} X_1 + S_1 \qquad S_1 \xrightarrow{\delta} \emptyset$$
$$\emptyset \xrightarrow{w_0^1} S_1$$
$$X_2 \xrightarrow{w_2^1} X_2 + A_1 \qquad A_1 \xrightarrow{\delta} \emptyset$$
$$S_1 + A_1 \xrightarrow{\gamma_1} \emptyset \qquad S_1 + C \xrightarrow{\gamma_2} C_1$$

Next, we write down the chemical reactions of node 2:

$$X_1 \xrightarrow{w_1^2} X_1 + S_2 \qquad S_2 \xrightarrow{\delta} \emptyset$$
$$X_2 \xrightarrow{w_2^2} X_2 + S_2$$
$$\emptyset \xrightarrow{w_0^2} A_2 \qquad A_2 \xrightarrow{\delta} \emptyset$$
$$S_2 + A_2 \xrightarrow{\gamma_1} \emptyset \qquad S_2 + C \xrightarrow{\gamma_2} C_2$$

Using the law of mass action, we can derive the ODEs:

$$\dot{s}_1 = w_0^1 + w_1^1 x_1 - \delta s_1 - \gamma_1 s_1 a_1 - \gamma_2 s_1 c \tag{50}$$
$$\dot{a}_1 = w_2^1 x_2 - \delta a_1 - \gamma_1 s_1 a_1 \tag{51}$$
$$\dot{s}_2 = w_1^2 x_1 + w_2^2 x_2 - \delta s_2 - \gamma_1 s_2 a_2 - \gamma_2 s_2 c \tag{52}$$
$$\dot{a}_2 = w_0^2 - \delta a_2 - \gamma_1 s_2 a_2 \tag{53}$$
$$\dot{c}_1 = \gamma_2 s_1 c - \delta c_1 \tag{54}$$
$$\dot{c}_2 = \gamma_2 s_2 c - \delta c_2. \tag{55}$$

The total amount of $c$ is constant as a result we can write down the mass conservation of $c$ as $c_1+c_2+c = c^{tot}$.

#### 2.2 Three sigma-based networks for dual region classifier

We begin with the chemical reaction of the first node::

$$X_1 \xrightarrow{w_1^1} X_1 + S_1 \qquad S_1 \xrightarrow{\delta} \emptyset$$
$$X_2 \xrightarrow{w_2^1} X_2 + S_1$$
$$\emptyset \xrightarrow{w_0^1} A_1 \qquad A_1 \xrightarrow{\delta} \emptyset$$
$$S_1 + A_1 \xrightarrow{\gamma_1} \emptyset \qquad S_1 + C \xrightarrow{\gamma_2} C_1$$

Next, we write down the chemical reactions of node 2:

$$\emptyset \xrightarrow{w_0^2} S_2 \qquad S_2 \xrightarrow{\delta} \emptyset$$
$$X_1 \xrightarrow{w_1^2} X_1 + A_2 \qquad A_2 \xrightarrow{\delta} \emptyset$$
$$X_2 \xrightarrow{w_2^2} X_2 + A_2$$
$$S_2 + A_2 \xrightarrow{\gamma_1} \emptyset \qquad S_2 + C \xrightarrow{\gamma_2} C_2$$

Next, we write down the chemical reactions of node 3:

$$C_1 \xrightarrow{w_1^3} C_1 + S_3 \qquad S_3 \xrightarrow{\delta} \emptyset$$
$$C_2 \xrightarrow{w_2^3} C_2 + S_3$$
$$\emptyset \xrightarrow{w_0^3} +A_3 \qquad A_3 \xrightarrow{\delta} \emptyset$$
$$S_3 + A_3 \xrightarrow{\gamma_1} \emptyset \qquad S_3 + C \xrightarrow{\gamma_2} C_3$$

Using the law of mass action, we can derive the ODEs:

$$\dot{s}_1 = w_1^1 x_1 + w_2^1 x_2 - \delta s_1 - \gamma_1 s_1 a_1 - \gamma_2 s_1 c \quad (56)$$
$$\dot{a}_1 = w_0^1 - \delta a_1 - \gamma_1 s_1 a_1 \quad (57)$$
$$\dot{s}_2 = w_0^2 - \delta s_2 - \gamma_1 s_2 a_2 - \gamma_2 s_2 c \quad (58)$$
$$\dot{a}_2 = w_1^2 x_1 + w_2^2 x_2 - \delta a_2 - \gamma_1 s_2 a_2 \quad (59)$$
$$\dot{s}_3 = w_1^3 c_1 + w_2^3 c_2 - \delta s_3 - \gamma_1 s_3 a_3 - \gamma_2 s_3 c \quad (60)$$
$$\dot{a}_3 = w_0^3 - \delta a_3 - \gamma_1 s_2 a_2 \quad (61)$$
$$\dot{c}_1 = \gamma_2 s_1 c - \delta c_1 \quad (62)$$
$$\dot{c}_2 = \gamma_2 s_2 c - \delta c_2 \quad (63)$$
$$\dot{c}_3 = \gamma_2 s_3 c - \delta c_3. \quad (64)$$

The total amount of $c$ is constant as a result we can write down the mass conservation of $c$ as $c_1 + c_2 + c_3 + c = c^{tot}$.

#### 2.3 Four sigma-based networks for non-linear classifier

We begin with the chemical reaction of the first node::

$$\emptyset \xrightarrow{w_0^1} S_1 \qquad S_1 \xrightarrow{\delta} \emptyset$$
$$X_1 \xrightarrow{w_1^1} X_1 + A_1 \qquad A_1 \xrightarrow{\delta} \emptyset$$
$$X_2 \xrightarrow{w_2^1} X_2 + A_1$$
$$S_1 + A_1 \xrightarrow{\gamma_1} \emptyset \qquad S_1 + C \xrightarrow{\gamma_2} C_1$$

Next, we write down the chemical reactions of node 2:

$$X_1 \xrightarrow{w_1^2} X_1 + S_2 \qquad S_2 \xrightarrow{\delta} \emptyset$$
$$X_2 \xrightarrow{w_1^2} X_2 + A_2 \qquad A_2 \xrightarrow{\delta} \emptyset$$
$$\emptyset \xrightarrow{w_0^2} A_2$$
$$S_2 + A_2 \xrightarrow{\gamma_1} \emptyset \qquad S_2 + C \xrightarrow{\gamma_2} C_2$$

Next, we write down the chemical reactions of node 3:

$$X_2 \xrightarrow{w_2^3} X_2 + S_3 \qquad S_3 \xrightarrow{\delta} \emptyset$$
$$X_1 \xrightarrow{w_1^3} X_1 + A_3 \qquad A_3 \xrightarrow{\delta} \emptyset$$
$$\emptyset \xrightarrow{w_0^3} A_3$$
$$S_3 + A_3 \xrightarrow{\gamma_1} \emptyset \qquad S_3 + C \xrightarrow{\gamma_2} C_3$$

Next, we write down the chemical reactions of node 4:

$$
\begin{array}{ll}
\emptyset \xrightarrow{w_0^4} S_4 & S_4 \xrightarrow{\delta} \emptyset \\
C_1 \xrightarrow{w_1^4} C_1 + A_4 & C_2 \xrightarrow{{}^4_2} C_2 + A_4 \\
C_3 \xrightarrow{w_3^4} C_3 + A_4 & A_4 \xrightarrow{\delta} \emptyset \\
S_4 + A_4 \xrightarrow{\gamma_1} \emptyset & S_4 + C \xrightarrow{\gamma_2} C_4
\end{array}
$$

Using the law of mass action, we can derive the ODEs:

$$\dot{s}_1 = w_0^1 - \delta s_1 - \gamma_1 s_1 a_1 - \gamma_2 s_1 c \tag{65}$$
$$\dot{a}_1 = w_1^1 x_1 + w_2^1 x_2 - \delta a_1 - \gamma_1 s_1 a_1 \tag{66}$$
$$\dot{s}_2 = w_1^2 x_1 - \delta s_2 - \gamma_1 s_2 a_2 - \gamma_2 s_2 c \tag{67}$$
$$\dot{a}_2 = w_0^2 + w_2^2 x_2 - \delta a_2 - \gamma_1 s_2 a_2 \tag{68}$$
$$\dot{s}_3 = w_2^3 x_2 - \delta s_3 - \gamma_1 s_3 a_3 - \gamma_2 s_3 c \tag{69}$$
$$\dot{a}_3 = w_1^3 x_1 + w_0^3 - \delta a_3 - \gamma_1 s_2 a_2 \tag{70}$$
$$\dot{s}_4 = w_0^4 - \delta s_4 - \gamma_1 s_4 a_4 - \gamma_2 s_4 c \tag{71}$$
$$\dot{a}_4 = w_1^4 c_1 + w_2^4 c_2 + w_3^4 c_3 - \delta a_4 - \gamma_1 s_4 a_4 \tag{72}$$
$$\dot{c}_1 = \gamma_2 s_1 c - \delta c_1 \tag{73}$$
$$\dot{c}_2 = \gamma_2 s_2 c - \delta c_2 \tag{74}$$
$$\dot{c}_3 = \gamma_2 s_3 c - \delta c_3 \tag{75}$$
$$\dot{c}_4 = \gamma_2 s_4 c - \delta c_4 \tag{76}$$
$$\tag{77}$$

The total amount of $c$ is constant as a result we can write down the mass conservation of $c$ as $c_1 + c_2 + c_3 + c_4 + c = c^{tot}$.

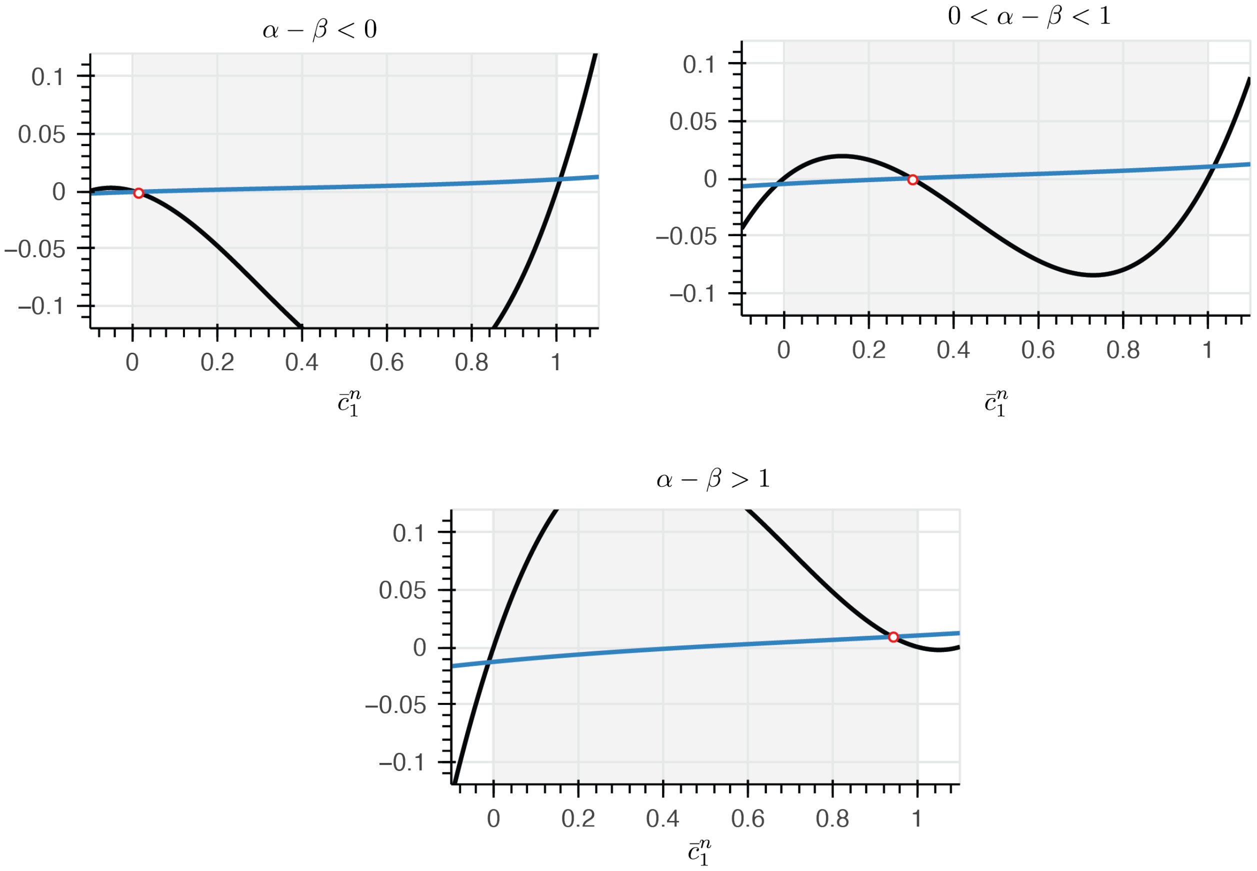

***Figure S1:*** **Plots analyzing the roots of equation** (22) **as a function of** $\alpha - \beta$.

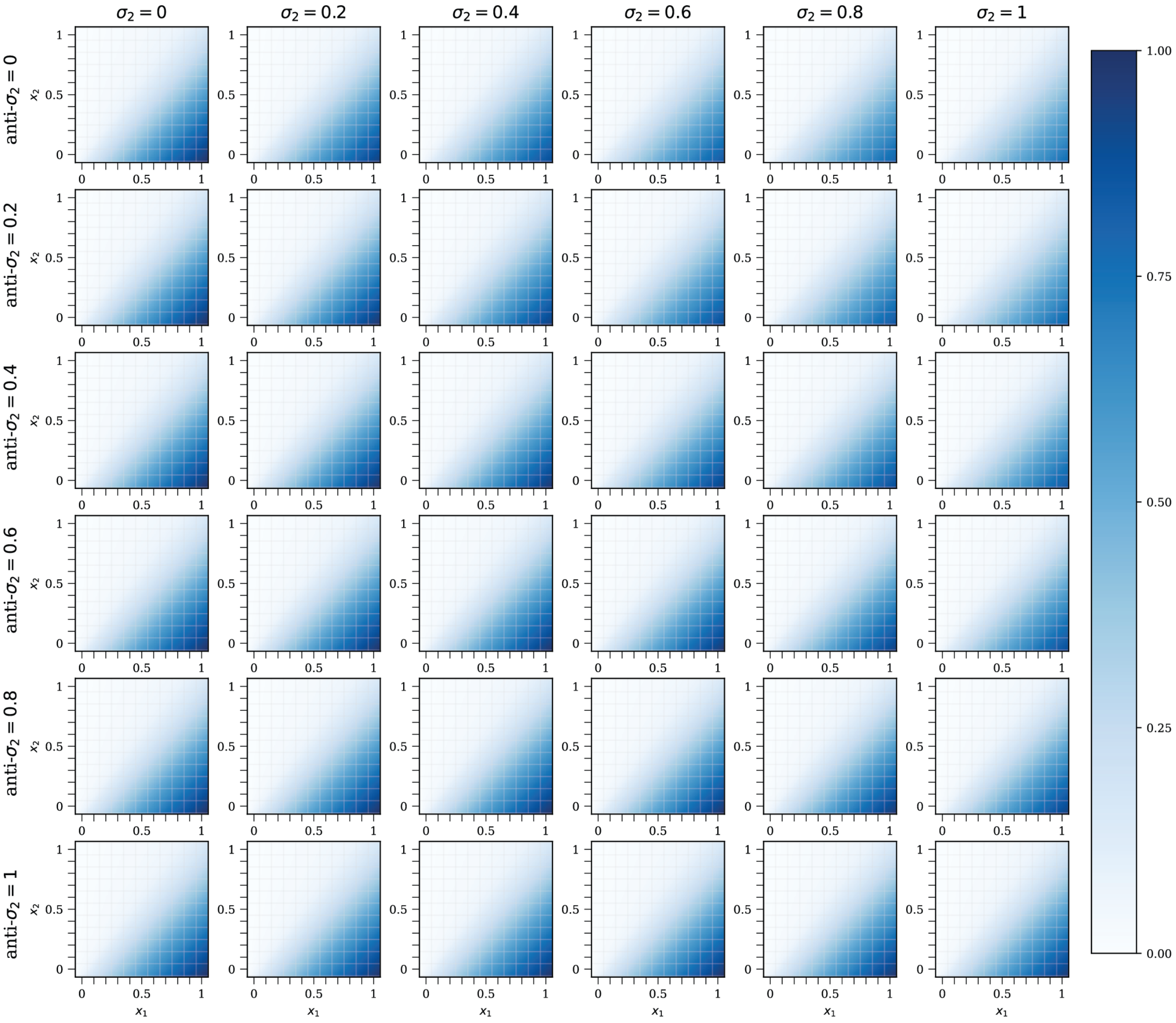

***Figure S2:*** **The effect of competition imposed by sigma 2 ($S_2$) and its corresponding anti-sigma ($A_2$) over a wide range of concentrations on perceptron output**. *Shown is a matrix of responses of the perceptron made by sequestration relationship of $S_1$ and $A_1$ over a wide range of concentrations of competing perceptron made of $S_2$ and $A_2$. In all simulations, $r = 0.01$. Higher $S_2$ results in attenuation of the response amplitude while leaving the pattern intact. Increasing $A_2$ sequesters $S_2$, thus leaving more resources for the $S_1$ which reveals itself in the recovery of response amplitude.*

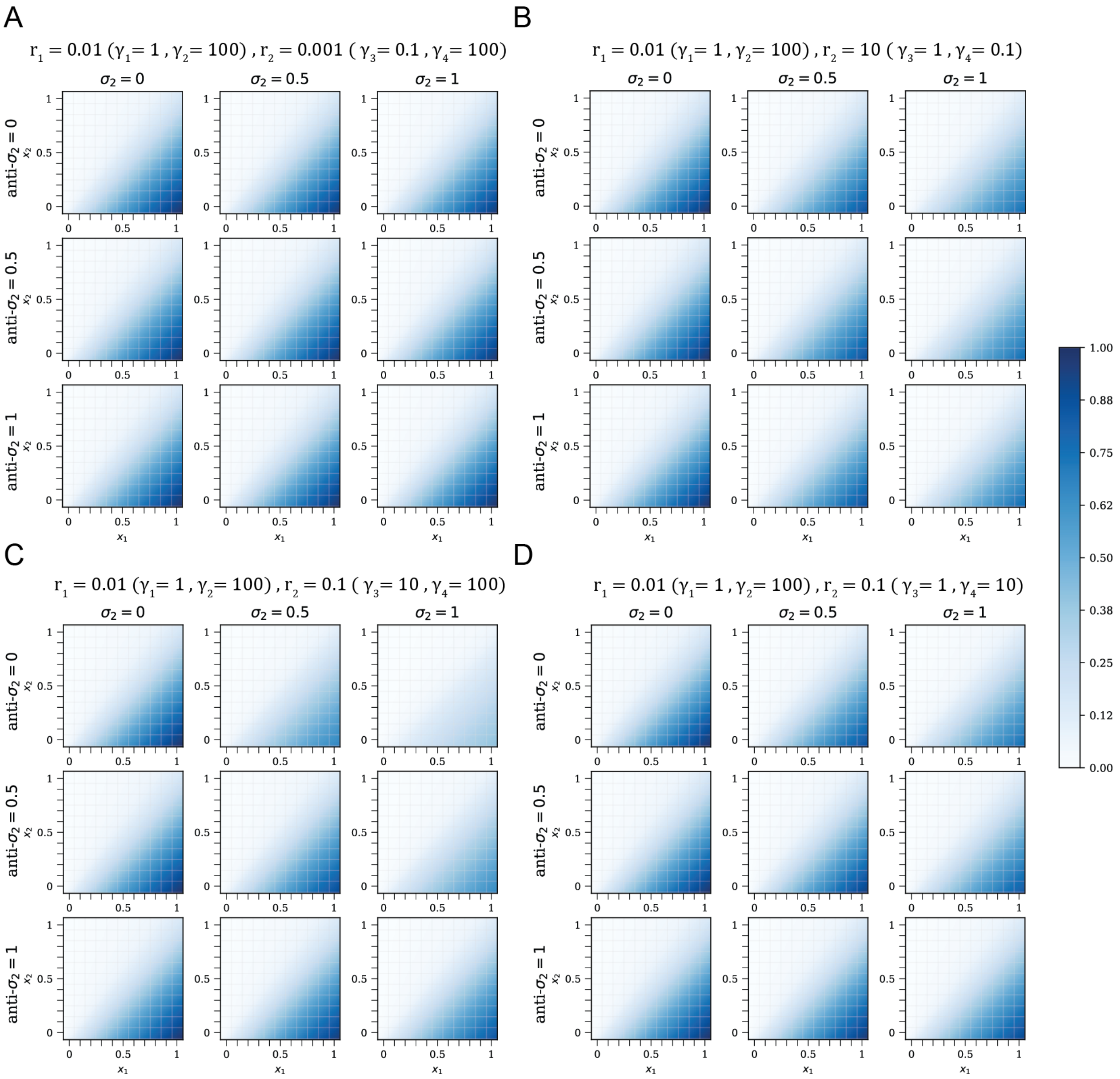

***Figure S3:*** **The kinetics of competing node only changes the amplitude of perceptron output**. *The response of a sequestration-based perceptron made of $S_1$-$A_1$ interaction is shown where the competitive binding ratio of $S_1$-$A_1$ denoted as $r_1$ is $0.01$ while $r_2$ is equal to $0.001$ in* ***A****, $0.01$ in* ***B****, $0.1$ in* ***C****, and $0.1$ in* ***D****. The affinity of $S_2$ binding with $C$ is denoted as $\gamma_3$ while the sequestration rate is shown as $\gamma_4$. The kinetics of $S_2$-$A_2$ binding which dictates the response of the competing perceptron merely tunes the amplitude of $S_1$-$A_1$ perceptron. When $\gamma_3$ is high (such as* ***C****), the competing perceptron consumes a majority of the resources, thus significantly suppressing the response in the absence of any sequestrating molecule ($A_2$). Similarly, when $\gamma_4$ is low such as* ***B****, the competing perceptron consumes resources with minimal sequestration effect. Therefore, it effectively only shrinks the available resources for the $S_1$-$A_1$ perceptron which can be observed from consistent suppressed response across all concentrations of $A_2$ with high $S_2$. In cases like* ***A*** *and* ***D*** *where the sequestration rate is stronger than complex formation, ($r \rightarrow 0$), the competition effect is minimal when complex formation is slow (as in* ***A****) or slight suppression when complex formation is moderate (such as* ***D****).*

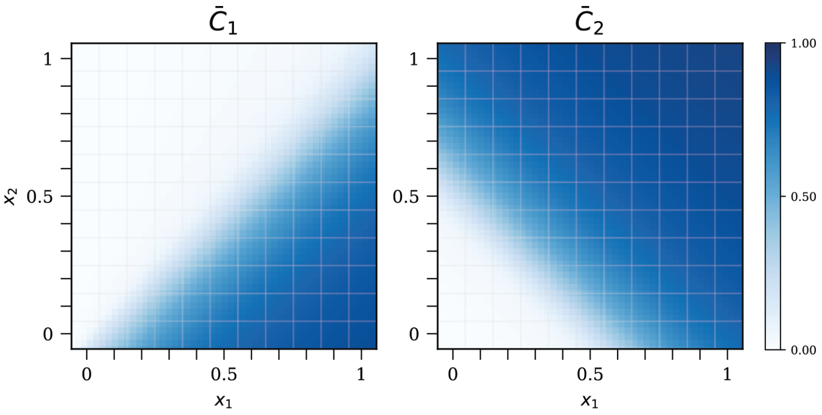

***Figure S4:*** **The decision boundary of two-perceptron network with shared inputs (Fig. 4A) in fast competitive binding regime ($r = 0.01$)**. *The decision boundary is linear since $r \ll 1$ and this linearized boundary directly affects the coupled network output (**Fig. 4**D)*

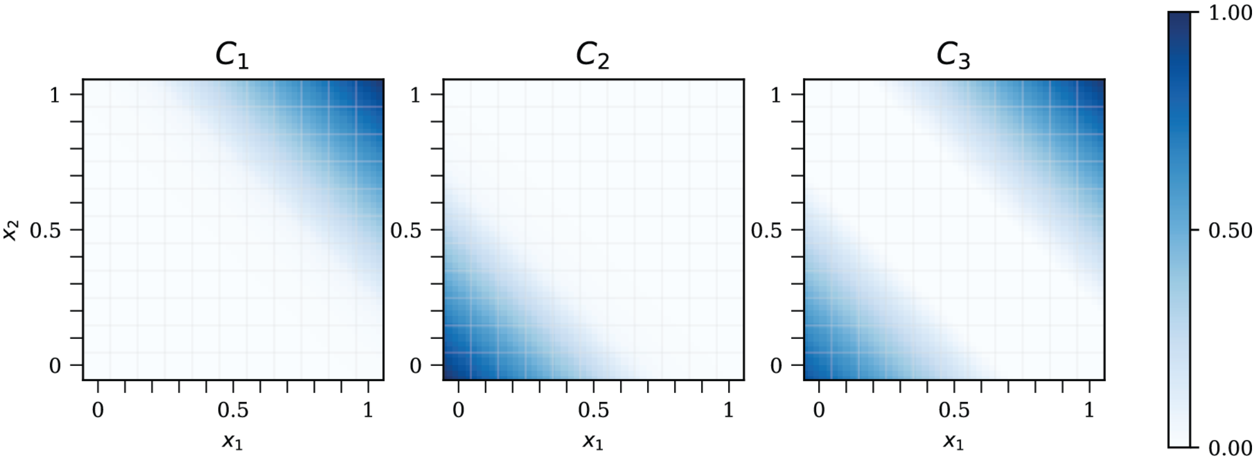

***Figure S5:*** **The ideal decision boundary of dual region classifier network in the absence of limited and shared resources**. *Since the first layer outputs $\bar{c}_1$ and $\bar{c}_2$ (left and middle, respectively) have no interference, the network output ($\bar{c}_3$, right) resembles the network response with shared resources **Figs. 4F** and* **G**

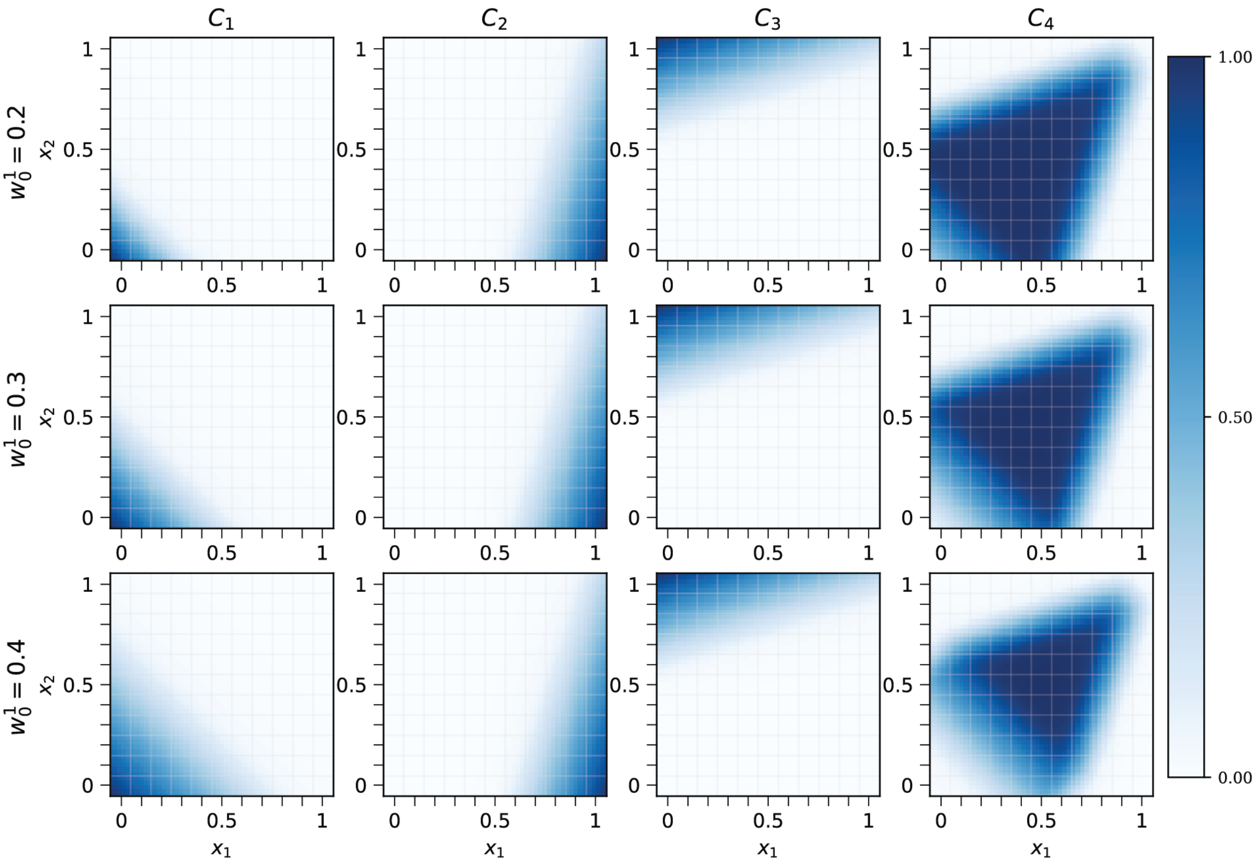

***Figure S6:*** **The ideal decision boundary of a band-pass network in the absence of limited and shared resources with different bias weights**. *The network output $\bar{c}_4$ is shown in the leftmost column while the first layer outputs ($\bar{c}_1$, $\bar{c}_2$, and $\bar{c}_3$) are shown in the other columns. Increasing the bias of the first node expands its decision boundary (left), thus affecting the non-linear decision boundary in the $[x_1, x_2] \to 0$ input regime.*

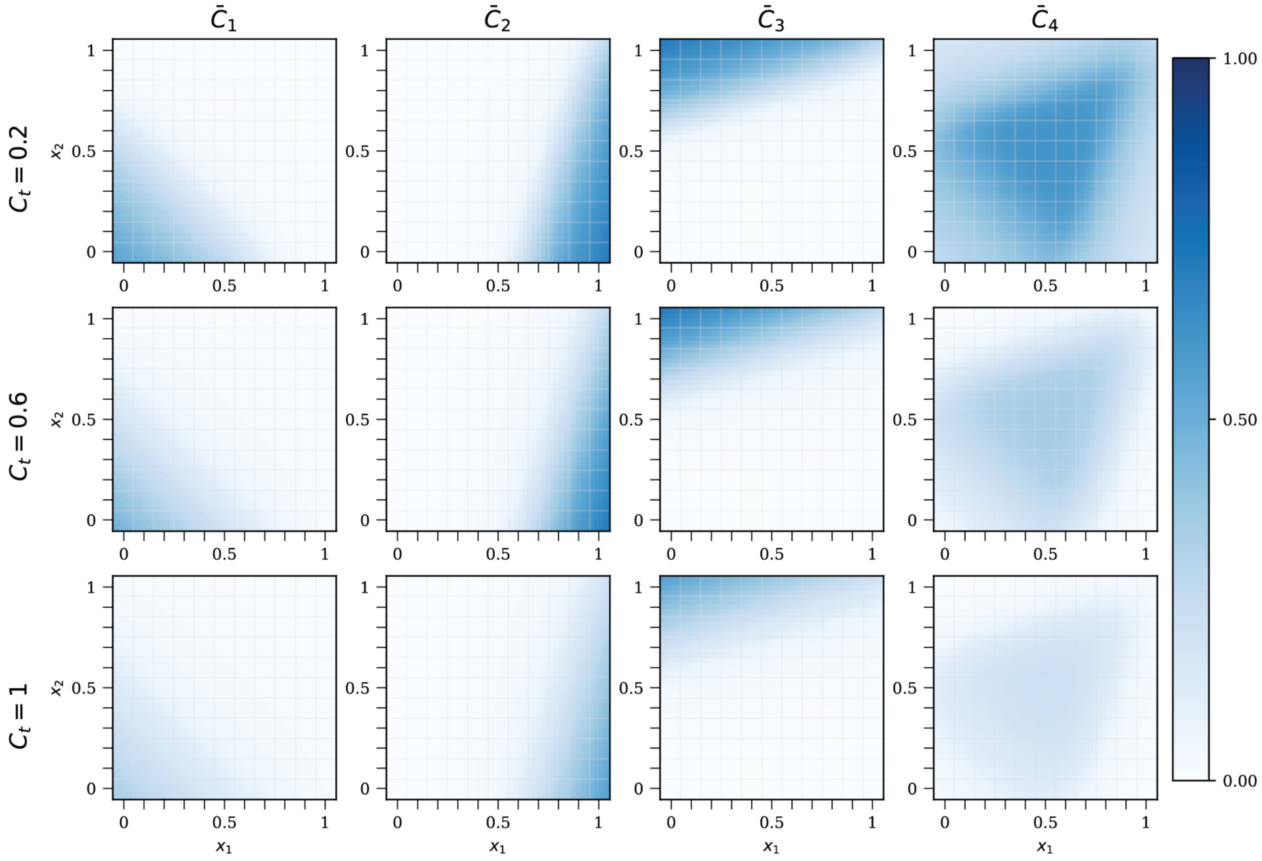

***Figure S7:*** **The non-linear classifier decision boundary in the presence of sharing various amounts of limited resources**. *The amplitude of the response $\bar{c}_4$ changes by increasing amounts of $c^{tot}$ while the response pattern remains mostly unchanged.*

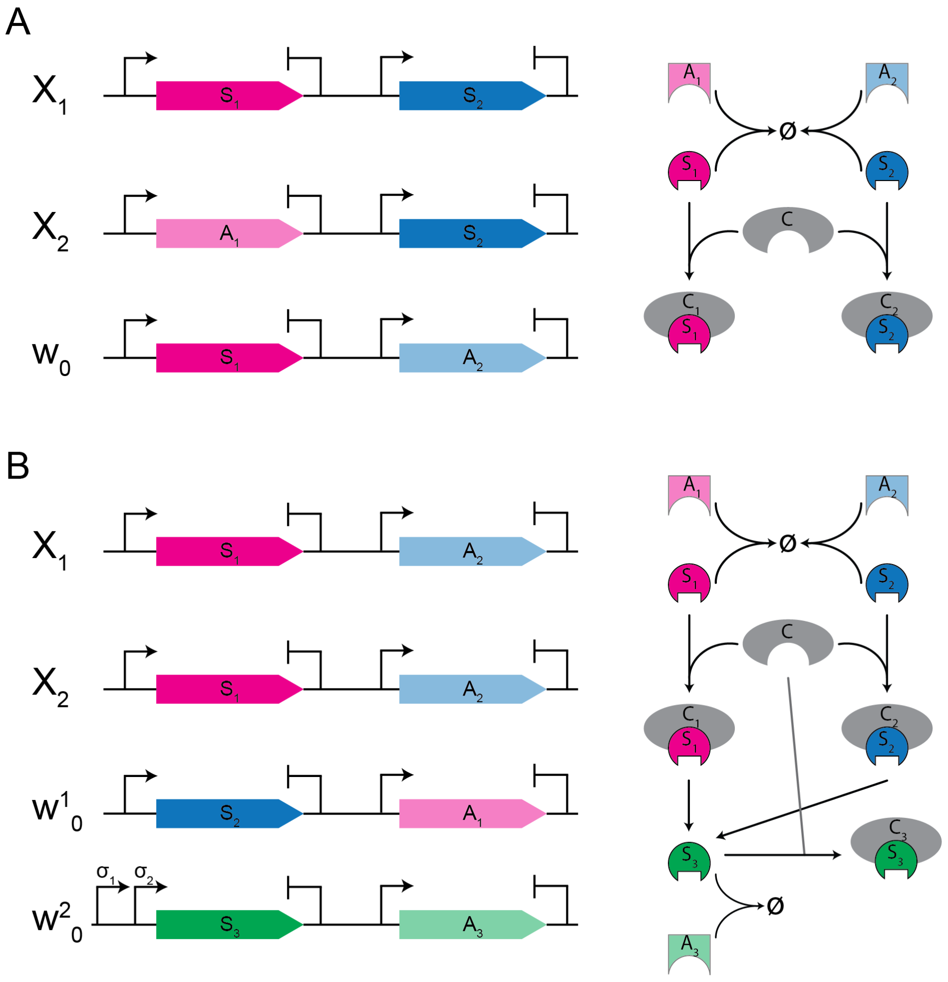

***Figure S8:*** **Schematic representation of DNA inputs required for reconstitution of non-linear neural networks presented in Figs. 4A and E in bacterial cells or cell-free expression systems.** *Promoter signs without a label stand for constitutive promoters and the strength of promoters and ribosome binding sites determine the weight of inputs.* ***A****, left: the schematic of DNA sequences required for construction of 2-perceptron network with shared inputs* ***Fig. 4A****. Varied copy numbers or concentrations of $X_1$ and $X_2$ plasmids determine the input of the network in a bacterial cell or cell-free system, respectively. Shown on the right is the schematic representation of the sequestration network.* ***B****, left: the schematic representation of DNA sequences that construct a dual region classifier* ***Fig. 4E****. Input bias $w_0^2$ encodes $S_3$ under promoters that bind to complexes $C_1$ and $C_2$. The sequestration interactions that generate the network are shown on the right.*

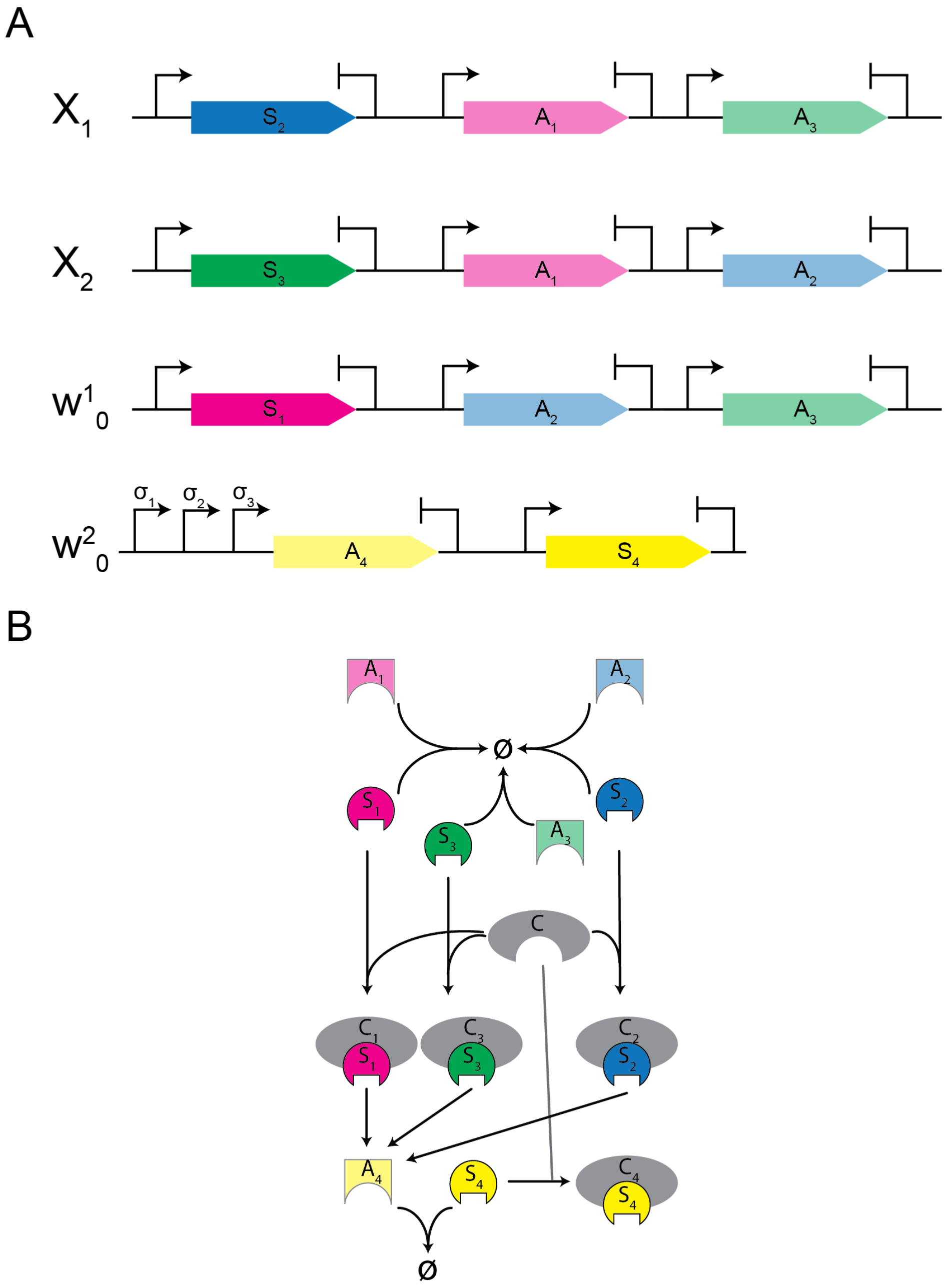

***Figure S9:*** **Schematic representation of DNA inputs required for construction of the non-linear classifier represented in Fig. 4H.** *Promoter signs without a label stand for constitutive promoters and the strength of promoters and ribosome binding sites determine the weight of inputs.* ***A****: the schematic representation of required DNA sequences to create a non-linear classifier (bandpass) in a bacterial cell or cell-free expression system. The input concentration can be controlled by modifying the plasmid copy number or its concentration in the cell-free expression reaction. Input bias $w_0^2$ encodes both $S_4$ and $A_4$ while controlling $A_4$ expression under the regulation of $C_1$, $C_2$, and $C_3$ complexes.* ***B****: the schematic representation of a network of sigma-anti sigma interactions that create the non-linear classifier presented in* ***Fig. 4H****.*
